# Learning interpretable representations of single-cell multi-omics data with multi-output Gaussian Processes

**DOI:** 10.1101/2024.11.03.621746

**Authors:** Zahra Moslehi, Florian Buettner

**Affiliations:** German Cancer Research Center (DKFZ), Heidelberg, Germany; Goethe University Frankfurt, Germany; German Cancer Consortium (DKTK), partner site Frankfurt/Mainz, a partnership between DKFZ and UCT Frankfurt-Marburg, Germany; Frankfurt Cancer Institute (FCI), Germany

## Abstract

Learning representations of single-cell genomics data is challenging due to the non-linear and often multi-modal nature of the data on one hand and the need for interpretable representations on the other hand. Existing approaches tend to either focus on interpretability aspects via linear matrix factorisation or on maximising expressive power via neural-network based embeddings using black-box variational autoencoders or graph embedding approaches. We address this trade-off between expressive power and interpretability by introducing a novel approach that combines highly expressive representation learning via an embedding layer with an interpretable multi-output Gaussian processes within a unified framework. In our model, we learn distinct representations for samples (cells) and features (genes) from multi-modal single-cell data. We demonstrate that even a few interpretable latent dimensions can effectively capture the underlying structure of the data. Our model yields interpretable relationships between groups of cells and their associated marker genes: leveraging a gene relevance map, we establish connections between cell clusters (e.g. specific cell types) and feature clusters (e.g., marker genes for those specific cell types) within the learnt latent spaces of cells and features.

## 1 Introduction

The single-cell genomics field has recently seen the development of many new techniques measuring different kinds of biomolecular features at the single-cell level. These methods include chromatin accessibility [1], gDNA profiling [2], methylation [3], chromatin immunoprecipitation profiling [4], protein [5], and lipid composition [6]. Each of these methods generates different data modalities that inform on different aspects of biological processes.

To integrate single-cell data from different data modalities, various computational methods have been developed.

Current state-of-the-art multi-omics integration methods aim to learn a representation of cells that integrate information from all data modalities. Interpreting this new space poses a significant challenge. To relate the learnt space to genes and other features, typical workflows first cluster cells in the latent space and then characterise these clusters via a differential expression analysis. Analysis on the cluster level, however, only results in a coarse-grained interpretability, omitting the structured varibility within clusters [7].

Rather than conducting such post-hoc differential expression analysis on the cluster level, we propose to explicitly model the inherent correlations and dependencies between samples and genes or other features within the data.

In this paper, we introduce **M**ulti-**O**mics **M**ulti-**O**utput **G**aussian **P**rocesses (MOMO-GP) for the integration of multi-omics data. MOMO-GP embeds samples, and features from different modalities (like genes and peaks) into separate interpretable latent representations. Using these representations for cells, genes, and peaks, along with gene relevance maps [8, 9] and peak relevance maps, we can directly encode cell-gene, and cell-peak relations. A group of cells can be related to a group of genes and a group of peaks. Learning these three embeddings jointly helps to achieve a high expressive power for each of the embeddings, whilst maintaining interpretability. MOMO-GP is not restricted to just genes or peaks and it can be used for any other view. Since MOMO-GP directly links samples, and features from different modalities together via their respective embedding spaces, it facilitates clustering-free marker detection as well as the cluster-agnostic analysis of feature-feature interactions. The direct encoding of cell-feature or feature-feature relations has been previously proposed in SIMBA, where all features and samples are co-embedded into a common latent space [10]. SIMBA constructs a graph in which cells and features are represented as nodes, and relations between these entities are encoded as edges. Then, a graph embedding approach is utilized to embed all nodes into a common low-dimensional space. However, the feature embeddings learnt by SIMBA only tend to have a limited expressive power, which may stem the inherent restriction to a single shared latent space between all cells and all features. Our results show that learning separate representations for features and cells substantially outperforms SIMBA, offering better expressive power and more faithful representations.

The primary concept of MOMO-GP is to learn a latent variable model that explicilty models dependencies between samples, features, and views. Standard latent variable models only model dependencies between samples and their multi-view versions between samples and views. We extend the framework of Gaussian Process Latent Variable Models (GP-LVM) [11, 9], a probabilistic kernel PCA via GP regression, which treats all features as independent. To explicitly model the dependencies between features (genes), we introduce an additional kernel to model the covariance between features. We then connect this feature kernel with the standard sample kernel that models dependencies between cells, via the Kronecker Product. For modelling multi-view data, we introduce additional kernels to capture dependencies among features of each view. We employ the Manifold Relevance Determination (MRD) approach [12] to learn for each dimension of the cell embedding, whether it is a private dimension that is specific to an individual view or whether it is shared between views and jointly models variance in multiple views.

In summary, MOMO-GP

- ….simultaneously learns a feature embedding for every modality and a shared cell embedding.
- …is designed to find a trade-off between expressive power and interpretability, by explicitly linking non-linear dependencies between features and cells.
- …outperforms other existing methods. In the sample space, it performs similar to other baseline and existing algorithms but provides better interpretability. In the feature space, our model outperforms SIMBA, the only other baseline to simultaneously learn and link feature and cell embeddings.

## 2 METHODS AND MATERIALS

### 2.1 Background

#### 2.1.1 A brief review on Gaussian Processes

Gaussian Processes (GPs) are a type of probabilistic model that defines a distribution over functions [13]. In GPs, a function is conceptualized as an infinite-dimensional vector, where a prior distribution is established over a set of *N* instances of them. This prior distribution follows a Gaussian distribution parameterized by a mean and a covariance. The mean is typically assumed to be zero, while the covariance is determined by a function of the input space on which the process operates. The covariance quantifies the similarity between all pairs from the input space, which is modeled by the kernel function. By sampling from the GP prior distribution, when a pair of input data points are close together, their function values are highly correlated. Consequently, this process yields a smooth function over the input space. When the input space is regarded as a latent variable, it is referred to as the GP Latent Variable Model (GP-LVM) [14, 11].

A multi-output Gaussian Process (GP) is an extension of the traditional single-output GP to simultaneously predict multiple correlated outputs, leveraging shared information across different tasks via a coregionalization matrix [15]. To more efficiently model the relation among different outputs and allow for new outputs at test time, in the Latent Variable Multiple Output Gaussian Processes (LV-MOGP) the coregionalization matrix is replaced by a kernel matrix [16]. LVMOGP then infers a latent space representing the information about different outputs. The kernel of this multi-output GP is separable and can be expressed as the Kronecker product of two individual kernels. The first kernel captures the similarity between samples, in the input space, while the second kernel measures the similarity between pairs of features. Note that in contrast to the GP-LVM, inputs are observed.

#### 2.1.2 GP Latent Variable Model (GP-LVM)

Here, we briefly explain the mathematical foundation of Gaussian Process Latent Variable Models (GP-LVM). Let the observed data in a high-dimensional space be denoted by **Y** ∈ ℝ^*I×J*^, where *I* represents the number of samples and *J* denotes the number of features. The matrix **Y** is considered a noisy version of the true values **F**∈ ℝ^*I×J*^. The relationship between **Y** and **F** is described by a likelihood function. We define a nonlinear mapping between the high-dimensional data **F** and a lowdimensional latent representation 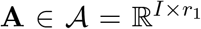, where *r*_1_ is the number of dimensions in the latent space, with *r*_1_ ≪ *J*. This mapping is governed by a Gaussian Process (GP), with **A** serving as the latent inputs.

The GP-LVM method assumes that the features are independent, while the samples exhibit strong correlations. The GP learns the correlation structure between the data points in the high-dimensional space by inferring **A** in a way that ensures a smooth mapping from the latent to the data space. This model maintains the integrity of dissimilarities, meaning that two points far apart in the data space cannot be positioned too closely in the latent space, as such proximity would imply a discontinuity in the mapping [17].

### 2.2 MOMO-GP algorithm

#### 2.2.1 Probabilistic model of MOMO-GP - single view version

In this section, we will briefly introduce the main concept of our probabilistic model, starting with the single-view version. The detailed formulation of the model is presented in the supplementary methods section.

In single-cell RNA-seq datasets, dependencies exist between different samples as well as different features. For example, cells of a specific cell type have a high similarity, and there are dependencies among all marker genes of similar cell types. Inspired by the LV-MOGP [16] and following [18], we model these dependencies between output dimensions via a kernel matrix and introduce an additional *r*_2_-dimensional latent variable 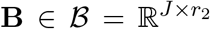 to the standard GP-LVM in order to model the correlation structure between the genes. Note that in LV-MOGP, the inputs are observed, whereas in our model, both latent variables **A** and **B** need to be inferred.

To facilitate an efficient implementation of the model, we follow [18] and represent the observed data via a triple store where an observed training sample is represented as (*i, j, y*_*i,j*_), where ∀ (*i, j*) ∈ [1, *I*] *×* [1, *J*] with sample *i*, feature *j* and corresponding entries in the observed matrix *y*_*i,j*_. In this way, the long vectors **y** ∈ ℝ^*I·J*^ is defined. Following the idea of LV-MOGP to define the dependencies of samples and features, a new coregionalization kernel needs to be defined as the Kronecker product of two individual kernels, one on the latent inputs and one on the latent outputs. Since this Kronecker product computes the correlation for all combinations of *I* samples and *J* features in matrix **F**, the size of the coregionalization kernel is (*I•J*) *×* (*I•J*).

We finally write our model as: *p*(**f**) = *𝒩* (**f** |**0, K**^coreg^), where **f** ∈ ℝ ^(*I•J*)^ and **K**^coreg^ ∈ ℝ ^(*I•J*)*×*(*I•J*)^ and the vector **y** ∈ ℝ^*I·J*^ is defined as the noisy version of **f**. To compute this GP model, we need to compute the inverse of the covariance matrix **K**^coreg^, which in a naive implementation has a complexity of 𝒪 (*n*^3^), where *n* is the number of samples, in this case *n* = *I•J*, where *I* is the number of samples and *J* is the number of features. In genomics data, we often have a large number of cells (*I*) as well as genes (*J*). To decrease the time complexity of the model and make the problem tractable, we employ the idea of sparse GPs. The fundamental idea behind sparse Gaussian Processes (GPs) is to approximate the full GP model using a smaller set of representative points known as inducing points [19]. These inducing points are significantly fewer than the original data points and effectively summarize the essential information in the data. They are selected to ensure that the GP model can be well-approximated with this smaller set, allowing the model to capture the core structure of the data without considering all data points simultaneously, thereby reducing computational complexity. For this purpose, we define the variables 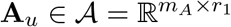, and 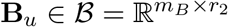. Here, **A**_*u*_ and **B**_*u*_ are inducing points in the latent spaces 𝒜 and ℬ, respectively. By leveraging this concept, we can already reduce the time complexity of our model from 𝒪((*I•J*))^3^ to 𝒪((*I•J*) *•* (*m*_*A*_*•m*_*B*_)^2^),where *m*_*A*_ and *m*_*B*_ are a subset of samples and features (inducing points) with *m*_*A*_ ≪ *I* and *m*_*B*_ ≪ *J*. Moreover, we enforce the same number of inducing points *m* for **A**_*u*_ and **B**_*u*_ and that allows us to replace the Kronecker product with an elementwise product, as proposed in [18]. Using this trick, we can then further reduce computational complexity to 𝒪((*I •J*) *m*^2^)). We empirically confirm this linear complexity for up to 7,000,000 entities in Figure S11.

In our model, the variables that need to be optimized include **A, A**_*u*_, **B, B**_*u*_, and other kernel parameters. To capture the nonlinear structure of the data, we follow the approach proposed in [18] and combine an embedding layer with a Gaussian Process layer. Instead of directly optimizing the variables **A** and **B**, we use an embedding function that embeds each cell and each feature into dense vectors of fixed size.

More formally, we map all indices in the range 1, …, *I* (representing cells) and 1, …, *J* (representing features) to matrices of size *I* × *r*_1_ and *J* × *r*_2_, respectively, using an embedding layer. Here, *r*_1_ represents the size of the input embedding space, and *r*_2_ represents the size of the output embedding space. For computing **A**_*u*_ and **B**_*u*_, we randomly select from 1, …, *I* and 1, …, *J*, respectively, and pass them through the embedding layer to obtain matrices **A**_*u*_ and **B**_*u*_ of size *m* × *r*_1_ and *m* × *r*_2_, respectively. During training, the weights of this embedding layer are optimized.

Figure 1 provides a graphical illustration of this proposed probability model. In this graphical model, shaded and white nodes represent observed and latent variables, respectively, while black circles denote parameters that need to be optimized by deriving the likelihood function. For further information, please refer to the supplementary Section. Additionally, Algorithm 1 in the supplementary notes outlines all the steps involved in the implementation process.

**Figure 1:**
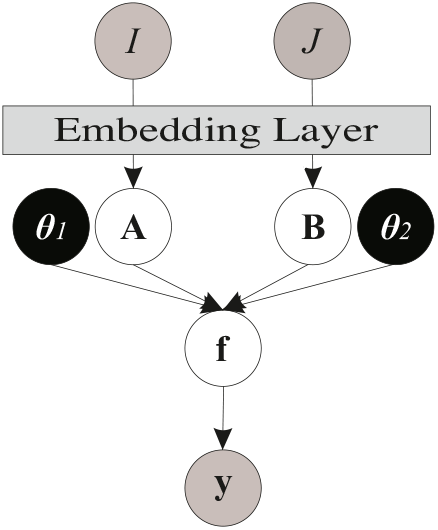
Probabilistic graphical model for single-view version of MOMO-GP. In this model, *θ*_1_ and *θ*_2_ are the parameter of the covariance functions.

#### 2.2.2 Probabilistic model of multi-view MOMO-GP

In the multi-view version of our method, we integrate a latent variable model that considers dependencies between samples, features, and views. This is different from standard latent variable models, which typically only model dependencies between samples or between samples and views. Similar to the single-view version, we use a kernel to model the covariance between samples. Then, we introduce additional kernels to capture dependencies among features of each view. Since the samples between different modalities are shared, we use one embedding for the samples but learn indivdual feature representations for each modality. Then, we link modalities via Manifold Relevance Determination (MRD) [12], which aims to decompose the representation of all data views into shared and private latent spaces. In brief, using Automatic Relevance Determination (ARD) priors [13], each view of the data is allowed to estimate a separate vector of ARD parameters. This view-wise relevance parameter allows us to learn for each dimension of the cell embedding, whether it is a private dimension that is specific to an individual modality or whether it is shared between modalities and explains variance in multiple modalities.

To illustrate how MOMO-GP can be extended to a two-view version, we consider single-cell gene expression data and single-cell ATAC-seq data.

Let *I* denote the number of samples, *J* denote the number of features for the first dataset (genes), and *K* denote the number of features for the second dataset (peaks). **Y**_1_ ∈ ℝ^*I×J*^ and **Y**_2_ ∈ ℝ^*I×K*^ represent our observed datasets generated from **F**_1_ and **F**_2_, respectively. However the same as single view version, we use a triple store for the both datasets. In this way, the long vectors **y**_1_ ∈ ℝ^*I•J*^ and **y**_2_ ∈ ℝ^*I•K*^ are defined. These are the noisy versions of **f**_1_ ∈ ℝ^*I•J*^ and **f**_2_ ∈ ℝ^*I•K*^. 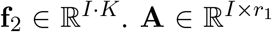 represents the low-dimensional embedding of cells, 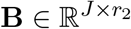 represents the embedding of genes, and 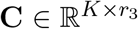 represents the embedding of peaks.

Rather than directly learning **A, B**, and **C**, we utilize an embedding layer to map row and column indices of the given datasets into these latent variables. We define one coregionalization kernel of size (*I •J*) *×* (*I•J*), formed by the Kronecker product of the covariance matrices **K**^**𝒜**^ and **K**^**ℬ**^, and another one of size (*I •K*) *×* (*I •K*), formed by the Kronecker product of **K**^**𝒜**^ and **K**^**𝒞**^. We then define two Gaussian processes, one for generating **f**_1_ using the first kernel and another one for generating **f**_2_ using the second kernel.

To fit the model, we define variables **A**_*u*_, and **B**_*u*_ to make the first Gaussian process sparse via inducing points, and **A**_*u*_, **C**_*u*_ are defined similarly for the second Gaussian process. Similar to **A, B**, and **C**, the variables **A**_*u*_, **B**_*u*_, and **C**_*u*_ are selected from the spaces 𝒜,ℬ, and 𝒞, respectively, but their sizes are much smaller than **A, B**, and **C**. We choose the same number of inducing points **A**_*u*_, **B**_*u*_, and **C**_*u*_ as proposed in [18], and replace the Kronecker product with an elementwise product to further reduce the time complexity of the algorithm. For more details, refer to the supplementary section.

In this model, the embedding of samples **A** is shared for generating both datasets **y**_1_ and **y**_2_. We utilize the idea of manifold relevance determination to allocate some latent dimensions of **A** shared between both datasets and some dimensions which are private for each dataset. Specifically, our kernel of samples **K**^𝒜^ would be different for **y**_1_ and **y**_2_. 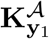 is an RBF kernel with Automatic Relevance Determination (ARD) of the form:

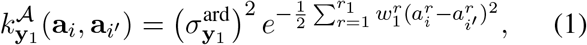

and similarly 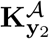 is defined. However, we learn a common latent space for both **y**_1_ and **y**_2_, but with the help of ARD weight vectors 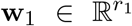 and 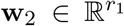, we can identify which dimensions are shared among both modalities and which dimensions are specifically assigned to each modality. In this way, the latent space **A** can be segmented as 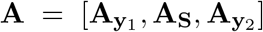, in which **A**_**S**_ is shared between both datasets for the set of dimensions *r* ∈ {1, …, *r*_1_} for which 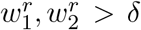, where *δ* is a number close to zero. The private space 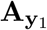 (resp. 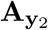) is defined for the set of dimensions for which 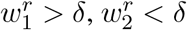 (resp. 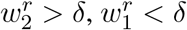).

Figure 2 graphically illustrates this model. The training algorithm of the multi-view version is similar to Algorithm 1 in supplementary section, but we consider that here we have two given datasets **y**_1_ and **y**_2_, two feature latent spaces **B** and **C**, and three sets of inducing variables, and thus two sets of **K**_*uu*_, **K**_*uf*_, and **K**_*fu*_ for generating **f**_1_ and **f**_2_. For optimizing **w**_1_ and **w**_2_, we update these ARD kernel parameters by maximizing the marginal likelihood distribution (13).

**Figure 2:**
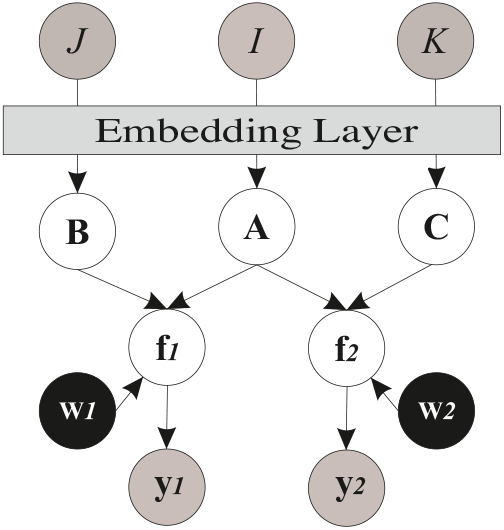
Probabilistic graphical model for multi view version of MOMO-GP.

#### 2.2.3 Implementation

The MOMO-GP model was implemented in Python using the GPFlow2 [20] and GPFlux [21] packages. The code for MOMO-GP is available at https://github.com/MLO-lab/MOMO-GP.git

### 2.3 Evaluation on single-cell data

#### 2.3.1 Single-cell RNA-seq and ATAC-seq integration

To evaluate our method, we utilized the PBMC 10k dataset from 10x Genomics, containing human Peripheral Blood Mononuclear Cells (PBMCs) from a healthy donor. This dataset encompasses paired single-cell multiome ATAC and gene expression sequencing. The dataset consists of 11,909 cells, 36,601 genes, and 134,726 peaks [22].

#### 2.3.2 CITE-seq integration

We also evaluated MOMO-GP on CITE-seq data of PBMCs. CITE-seq datasets contain transcriptome-wide measurements for single cells, including gene expression data and surface protein level information for a few dozen proteins. The dataset consists of 5247 cells, 33538 genes, and 32 proteins [23].

#### 2.3.3 Data pre-processing

For preprocessing, we utilized Scanpy [24] to perform normalization, logarithmic transformation, clustering, and cell type annotation for both single-cell RNA-seq (scRNA-seq) and single-cell ATAC-seq (scATAC-seq) datasets.

##### Single-cell RNA-seq

We applied quality control by filtering low-quality cells and those with high mitochondrial content. Genes detected in only a small number of cells were excluded. After normalization and logarithmic transformation, we used Leiden clustering to annotate cell types. Clusters showing noise, high ribosomal gene expression, or proliferating cells were removed. Further feature selection was performed to retain only the most variable and biologically relevant genes for downstream analysis.

##### Single-cell ATAC-seq

We filtered peaks detected in a minimal number of cells and retained cells with an appropriate number of accessible chromatin regions. Latent Semantic Indexing (LSI) was used for normalization, followed by the same lognormalization approach as in scRNA-seq. Clusters were annotated based on marker genes, and additional filtering focused on the most variable chromatin regions. Only cells passing the respective quality control criteria were retained in each modality. For integration purposes, only cells present in both modalities were considered.

The number of cells in the intersection of the RNA-seq and ATAC-seq datasets, used in our analysis of the PBMC 10k dataset, amounted to 9393.

##### Single-cell Protein Data

Protein expression data were normalized using the Denoised and Scaled by Background (DSB) method [25]. The dataset comprises 32 proteins, and the number of cells in the intersection of the RNA-seq and protein expression data, utilized in our analysis of the 5k PBMCs CITE-seq dataset, amounted to 3891.

### 2.4 Benchmarking

We compared our method with several commonly used baselines and related methods: Principal Component Analysis (PCA) [26], Uniform Manifold Approximation and Projection (UMAP) [27], Bayesian Gaussian Process Latent Variable Model (BGPLVM) [28], SCVI [29], and SIMBA [10].

PCA was selected as a linear dimensional reduction approach due to its ability to provide interpretable results. In our results, we ran PCA via Scanpy.

UMAP, a non-linear manifold learning algorithm widely used for visualizing biological data points, was also run via Scanpy. However, UMAP does not provide interpretable embeddings.

MOMO-GP extends the GP-LVM by incorporating dependencies for both inputs and outputs, while GP-LVM assumes all output values are independent. We evaluate a Bayesian implementation (BGPLVM) as a non-linear model that provides interpretable results but does not support feature embedding. We utilized the implementation of BGPLVM developed with GPytorch [30].

SCVI was selected as a state-of-the-art algorithm in neural network-based embedding algorithms. For SCVI, we employed the SCVI-tools package [29], which is designed for single-cell data and built on PyTorch and AnnData.

SIMBA is a method for co-embedding samples and features. To the best of our knowledge, it is the only method that learns and links both sample and feature embeddings in single-cell data. We compared our sample and feature embedded data with the outputs of SIMBA both qualitatively and using quantitative metrics.

### 2.5 Evaluating the quality of the results

For analyzing the results, we use two different quantitative metrics.

When cell type information is available, we can use accuracy as a criterion to evaluate the results. First, we apply clustering on the embedded data. To compute clustering accuracy, we assign the predicted label as the most frequent class label of each cluster to all of its data points. Then, accuracy is computed by dividing the total number of data points with the correct predicted label by the total number of all data points. Formally, it is as follows:

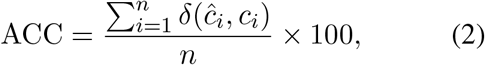

where *n* is the number of data points, *c*_*i*_ is the correct real label, *ĉ*_*i*_ is the predicted label, and the delta function *δ*(*s, t*) = 1 when *s* = *t*, otherwise it is 0 [31, 32].

The other metric used is the Adjusted Rand Index (ARI) [33]. ARI evaluates the similarity between two data clusterings. It considers all pairs of samples, counting those assigned to the same or different clusters in both the predicted and true clusterings. In our case, one of the clusterings would be the grouping of data points based on their cell types. This allows us to evaluate the discrimination of cell types provided by our learned embedding. Unlike the raw Rand Index, the Adjusted Rand Index (ARI) adjusts for the chance grouping of elements, providing a more accurate assessment of clustering performance. By adjusting for the chance grouping, the ARI provides a robust metric for comparing the similarity of cluster assignments.

In our experiments, we provide the ACC and ARI values for both sample embeddings and feature embeddings.

## 3 RESULTS

### 3.1 Single-cell RNA-seq analysis with MOMO-GP

While we propose a multi-omics algorithm with sample and feature embedding, we first verify the effectiveness of our method in the single omics case. Specifically, we check if our embeddings are competitive compared to popular algorithms. This initial validation ensures that our model can produce high-quality embeddings before extending its application to multi-omics data. To this end, we utilized the RNA modality of the PBMC 10k dataset to assess the performance of single-view MOMO-GP. Similarly, we used the RNA modality of the 5k PBMC CITE-seq dataset for evaluation purposes. The results are evaluated across various aspects:

#### 3.1.1 Cell Embedding

Our approach using multi-output Gaussian processes focuses on learning a low-dimensional embedding where each dimension is interpretable. Unlike linear methods, our non-linear approach allows us to use only a handful of latent variables (LVs) to model the data effectively. This results in a low-dimensional representation that maintains both interpretability and non-linearity, providing a faithful and meaningful representation of the underlying data structure. In this section, we demonstrate that the MOMO-GP embedding of cells is comparable to or better than other existing methods.

We projected the gene expression data points into a two-dimensional space after applying PCA [26], UMAP [27], and BGPLVM [28] algorithms. The results of the PBMC 10k dataset are depicted in Figure 3 and Supplementary Figure S1 (and Supplementary Figure S2 for the PBMC 5k-CITE-seq dataset). We ran SCVI [29] and our method in three different setups: embedding data points into a 32-dimensional space and displaying the 2D visualization of UMAP embedding of projected data; 2D embedding of data; and embedding data points into a 4D space and selecting the best results of 2 latent factors from these 4 different dimensions. The results for other 2 latent factors from these 4 different dimensions are given in Supplementary Figure S3. Data points in this figure are colored based on their cell type. After applying the MOMO-GP algorithm, the separation between the 13 different cell types, including CD4+ naïve T, CD8+ activated T, naïve B, intermediate monocytes, MAIT, mDC, CD14 monocytes, memory B, CD8+ naïve T, pDC, CD16 monocytes, CD4+ memory T, and NK cells, is well-defined, and the components appear well-coordinated. For a more quantitative comparison, ACC and ARI values for different methods are also presented in this figure.

**Figure 3:**
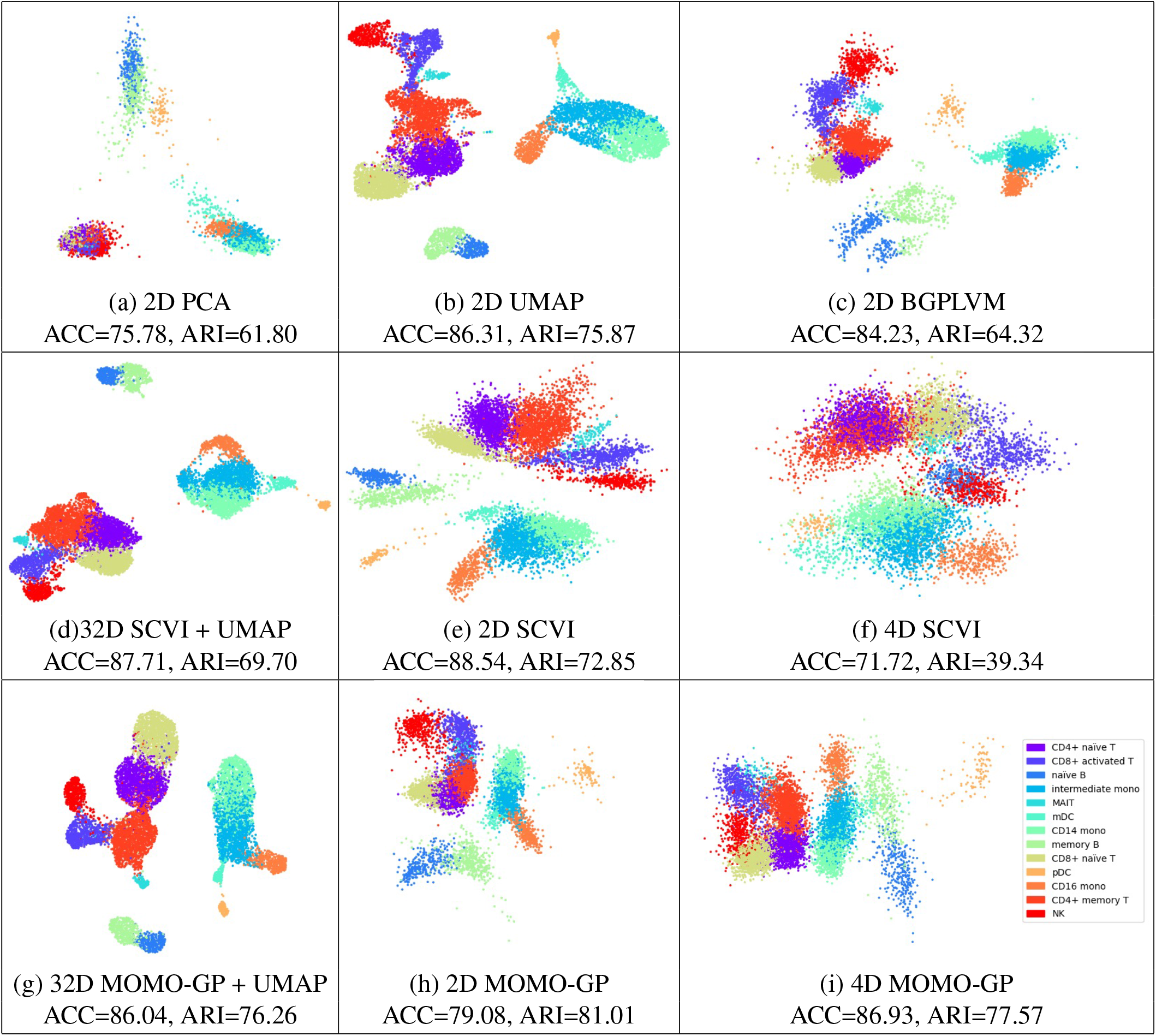
2D visualization of cells in the PBMC 10k dataset for scRNA-seq data using various methods: 2D PCA, (b) 2D UMAP, (c) 2D BGPLVM, (d) 32D SCVI+UMAP, (e) 2D SCVI, (f) 4D SCVI, (g) 32D MOMO-GP+UMAP, (h) 2D MOMO-GP, and (i) 4D MOMO-GP.

#### 3.1.2 Gene Embedding

In this section, we demonstrate that even with a few latent dimensions, the underlying structure of the data can be captured effectively without utilizing all genes. Specifically, we set the number of latent dimensions for both cell and gene representations to 2 and visualize the 2D embedding of all cells and genes. The results for the PBMC 10k dataset are depicted in Figure 4 (and Supplementary Figure S4 for the PBMC 5k-CITE-seq dataset). In these figures, cells are colored based on their cell types. Additionally, for each cell type, we identify the top 100 differentially expressed marker genes and color them according to their respective cell types. From the visualization, it is evident that our gene embedding using only 2 latent factors yields meaningful insights. Although there isn’t a perfect separation between marker genes of all different cell types, all marker genes of a specific cell type tend to form a cohesive cluster. Another interesting observation in this figure is the presence of a gray cluster in the middle of Figure 4.d. These genes do not exhibit specific biological associations with any particular cell types, leading them to form a distinct cluster within our embedding. To further elucidate the role of these genes, we selected the top 20 genes located near the center of the data, within the gray region (listed in Table S1). These genes are characterized by their involvement in diverse regulatory processes, including immune responses, development, and gene expression. Many of these genes are long non-coding RNAs (lncRNAs) involved in gene regulation (e.g. AC022445.1, EMX2OS, AC005481.1, AC024933.1, LINC02821, CARMN, AL590999.1, AC079035.1, AL589740.1, and AC092134.1).

**Figure 4:**
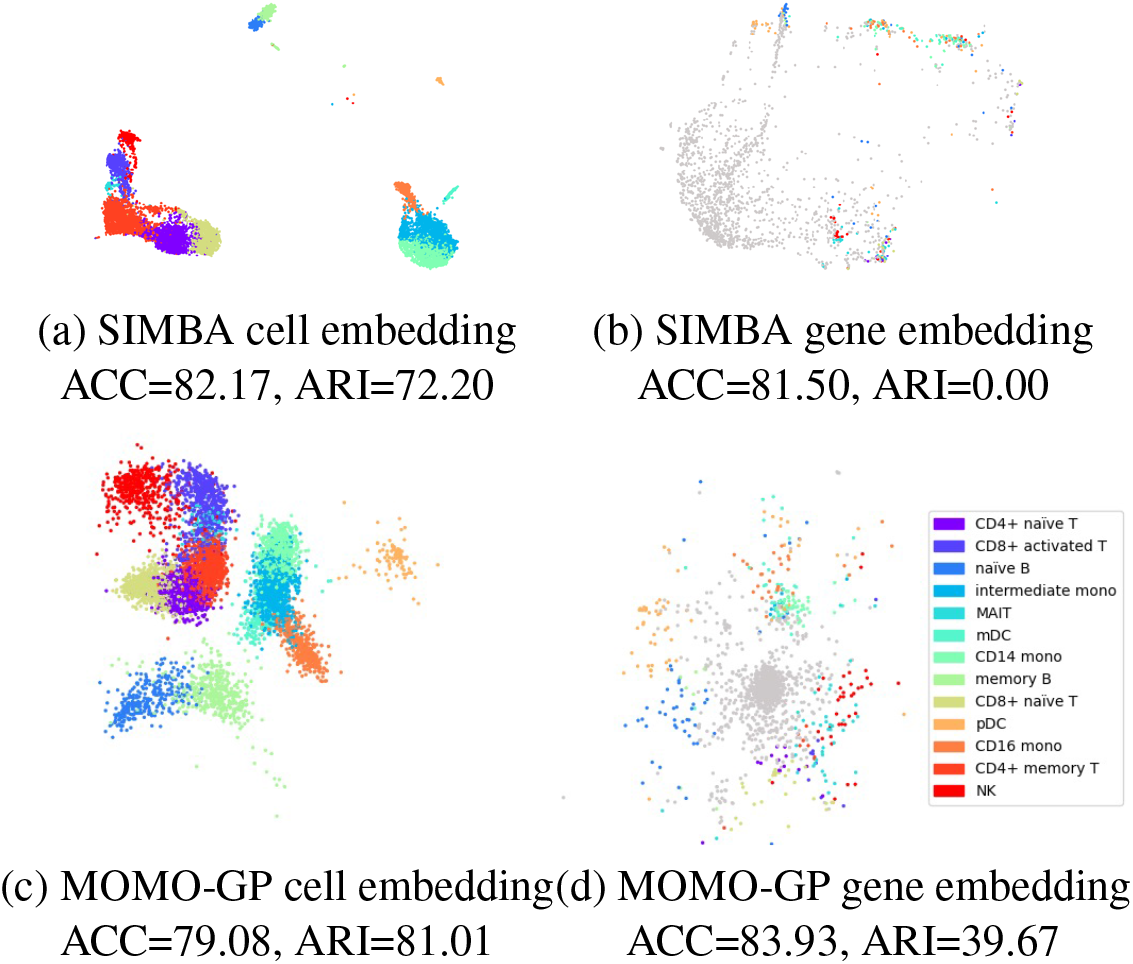
Visualization of PBMC 10k dataset using SIMBA and MOMO-GP embedding techniques for scRNA-seq data. (a) SIMBA-UMAP embedding of cells, with cell types color-coded, in a 50-D space. SIMBA-UMAP embedding of genes, highlighting the top 100 marker genes per cell type, color-coded by their respective cell types, in a 50-D space. Non-marker genes are shown in gray. (c) MOMO-GP embedding of cells in a 2D space. (d) MOMO-GP embedding of genes in a 2D space.

Additionally, we provide cell and gene embedding of RNA data from the PBMC 10k dataset using the SIMBA method. The default number of latent dimensions in SIMBA is set to 50. The cell and gene embeddings generated by SIMBA, followed by UMAP visualization, are presented in Figure 4.a and Figure 4.b, respectively. For the results of MOMO-GP with 50 latent dimensions followed by UMAP visualization, refer to Figure S5. While SIMBA’s cell embedding demonstrates effective separation among various cell types, its gene embedding noticeably underperforms compared to MOMO-GP.

#### 3.1.3 Interpretability of the Model

A significant characteristic of MOMO-GP is its capability to project both samples and features in a latent space. This feature becomes particularly valuable when we aim to establish connections between groups of samples and groups of genes in the latent space without relying on any ground truth about cell and gene labels. To achieve this, we adopt the concept of gene relevance maps [8, 9], the details of which are provided in the supplementary methods. In brief, a local gene relevance plot delineates the regions in a cell embedding where a gene’s contributions are most pronounced. In our analysis, instead of identifying the single highest relevant gene for each area, we opt to identify groups of meta-genes relevant to that area. We leverage the MOMO-GP gene embedding and identify meta-genes [7] (groups of similar genes) from our gene embedding. Subsequently, we link the highest globally relevant metagenes to certain cells using the concept of gene relevance maps. This approach enables us to link a group of genes (belonging to one meta-gene) to a group of cells. The outcomes of this experiment on the PBMC 10k dataset are depicted in Figure 5 (and Supplementary Figure S9 for the PBMC 5k-CITE-seq dataset). In Figure 5.a, we illustrate the gene embedding results, with all genes belonging to one meta-gene uniformly colored. In Figure 5.b, to show a “ground truth” about genes, we define the top 100 marker genes for each cell type and color them according to their corresponding cell type. For the cell embedding, we highlight the areas belonging to specific cell types by coloring all data points based on their cell type, as shown in Figure 5.c. These color codes will help us verify the cell types in Figure 5.c, check which genes belong to one meta-gene in Figure 5.a, and determine their relevant cell types by examining the marker genes in Figure 5.b. Figure 5.d delineates the areas where each meta-gene is relevant. To evaluate the results, we consider our knowledge about cell types in cell embedding and marker genes in gene embedding. For example, upon analyzing the gene relevance map for meta-gene 1, we observe that all cells in this area are CD8+Naïve T cells, and upon inspecting the genes of meta-gene 1, we note that most of them are marker genes of CD8+Naïve T cells. This process can be repeated for all meta-genes, enabling us to relate a group of cells to a group of genes using our embedding from MOMO-GP.

**Figure 5:**
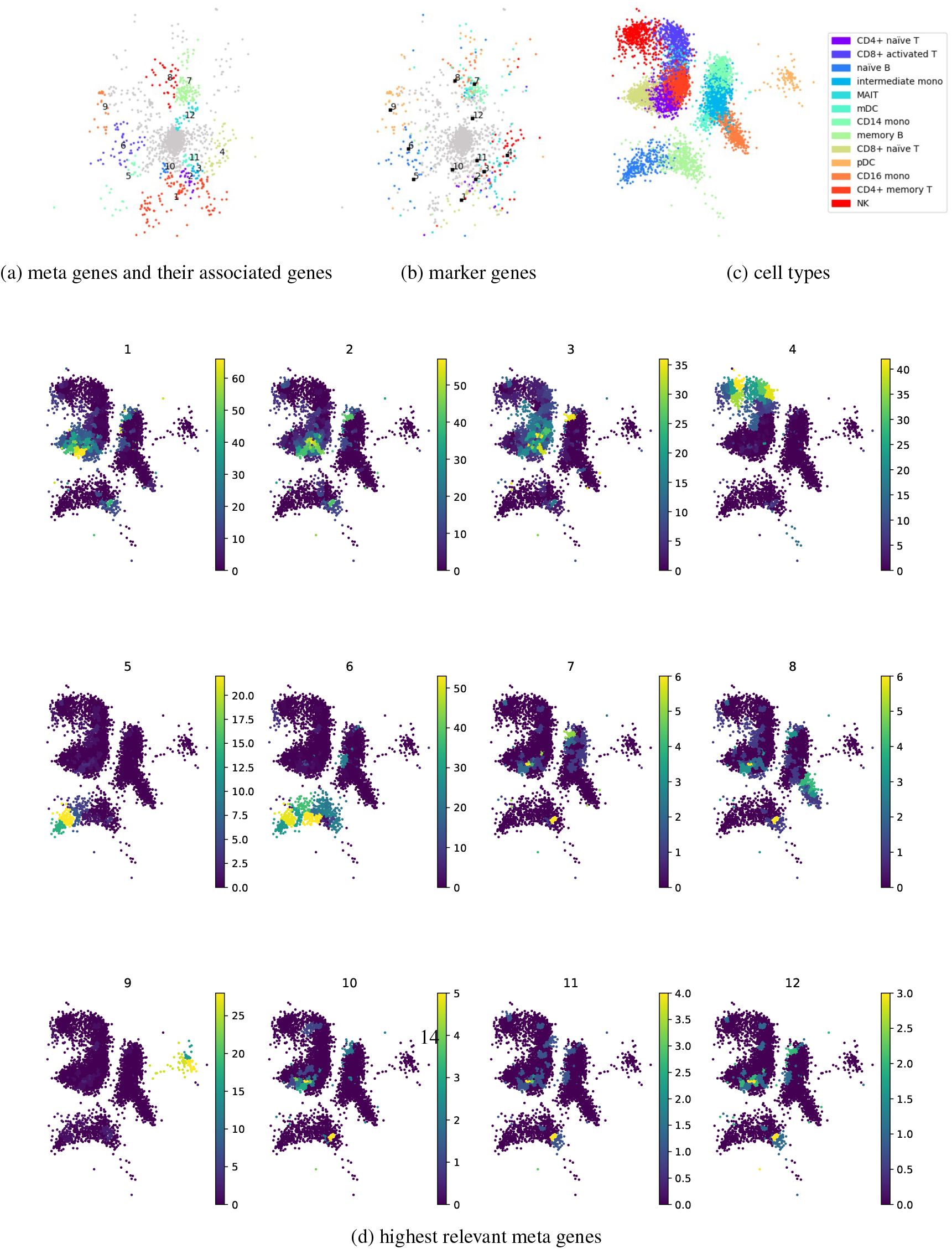
Exploration of the PBMC 10k dataset using a gene relevance map, which automatically identifies connections between groups of cells and genes: (a) Gene embedding colored according to genes associated with each meta-gene. (b) Gene embedding colored by marker genes specific to each cell type, (c) Cell embedding colored by cell types, (d) Gene relevance plot highlighting regions where gene contribution is highest. For instance, meta-gene 9 is enriched for pDC marker genes and exhibits significant relevance in the corresponding region of the cell embedding.

Furthermore, for additional evaluation, we would like to understand what the meta-genes are and whether they capture biologically meaningful gene sets. To do that, we employed gene set enrichment analysis [34] with Over-Representation Analysis (hypergeometric test) [35], implemented by the Gene Set Enrichment Analysis in Python (GSEAPY) package [36]. ORA aids in identifying gene sets that are predominantly present in our gene lists of interest. For this analysis, the gene lists comprise the genes of each meta-gene, while the gene set is selected from the human MSigDB collections [37]. Specifically, we select the C8 cell type signature gene set for bone marrow. The outcomes of this experiment on the PBMC 10k dataset are presented in Table 1 (and Supplementary Table S2 for the PBMC 5k-CITE-seq dataset). For each meta-gene, we sort enriched gene sets based on the combined enrichment score (computed with GSEAPY) and show the two most strongly enriched ones which adjusted p-value less than 0.05. Those meta-genes, that do not have any enriched gene sets are not shown in the table.

**Table 1:** PBMC 10K dataset: A list of gene sets enriched for each meta gene.

| Meta Gene | Term | Adjusted P-value | Combined Score | Cell-type Coverage |
| --- | --- | --- | --- | --- |
| 1 | Naïve T | 2.02e-43 | 2235.34 | 89.57 |
| 1 | CD8 T | 3.59e-02 | 26.99 | 49.16 |
| 2 | Naïve T | 2.73e-08 | 597.65 | 68.28 |
| 4 | NK | 8.38e-52 | 14262.21 | 34.71 |
| 4 | CD8 T | 3.46e-03 | 103.89 | 65.28 |
| 5 | Follicular B | 3.24e-06 | 391.46 | 100 |
| 6 | Follicular B | 6.53e-24 | 2545.37 | 100 |
| 6 | Plasma | 1.14e-03 | 107.62 | NA |
| 7 | Neutrophil | 1.17e-57 | 3013.32 | NA |
| 7 | Immature Neutrophil | 6.55e-21 | 786.68 | NA |
| 8 | Monocyte | 4.98e-23 | 4046.71 | 57.38 |
| 9 | Dendritic | 5.81e-20 | 4608.97 | 94.66 |
| 9 | CD34 B | 1.65e-02 | 109.18 | NA |

*Note:* To compute Cell-type Coverage for the term "Naïve T", we include all CD4+ Naïve T and CD8+ Naïve T cells. For the term "CD8 T", we count only CD8+ Naïve T cells. Due to overlaps between these two groups, the cumulative value for Meta Gene 1 exceeds 100 %.

Via the gene sets enriched in each meta-gene, we can approximately define the cell type associated to each group of genes. By comparing these enriched cell types with those relevant in the gene relevance map, we can validate first that MOMO-GP learns a meaningful gene embedding with similar genes being grouped together. Second, we validate that our relevance-based approach links gene- and cell-embeddings in a meaningful fashion. To do that, we have to identify the cells in which a meta-gene is relevant and check their cell types. Then, by comparing their cell types and the cell type associated to the respective meta-gene, we can validate that meta-genes capture meaningful groups of genes. For example, according to the ORA results of meta-gene 1, we observe a relationship between T-cells and genes of this meta-gene. On the other hand, via the gene relevance map, we find a relation between meta-gene 1 and T-cells. In Table 1, we present the cell type coverage values. For each meta-gene, we compute its gene relevance map and compute the fraction of cells with relevance score above a threshold *τ* that matches the cell type predicted by GSEA. For example, in the case of meta-gene 1, 89.57 percent of the cells with a relevance score above *τ* = 30 are classified as Naive T cells. These results highlight the strong structure within the gene-embedded data generated by MOMO-GP and its clear and meaningful relationship with cell embeddings.

### 3.2 Single-cell multi-omics integration with MOMO-GP

To demonstrate the effectiveness of our model on multi-view data, we examine its performance on two datasets: the PBMC 10k dataset, which combines paired scRNA-seq and scATAC-seq data, and the 5k PBMC CITE-seq dataset, comprising gene expression data and protein level information. We quantify the quality of cell embeddings and feature embeddings for all modalities.

#### 3.2.1 Cell Embedding

In this section, we illustrate the representation of cells using both our method and the SIMBA algorithm. We projected the paired scRNA-seq and scATAC-seq data from the PBMC 10k dataset, with our MOMO-GP mapping the data into a 2-dimensional space and SIMBA mapping the data into a 50-dimensional (its default value) space. Subsequently, we applied the UMAP method to visualize the results using SIMBA. The findings are depicted in Figure 6.a and 6.d. Embedding of cells into 50-dimensional space using MOMO-GP followed by UMAP is given in Figure S6.a. Our method demonstrates a notable separation between different cell types, with all samples for each cell type clustered together, while SIMBA produces separate clusters for each cell type. Similarly, the cell embedding of MOMO-GP on the PBMC 5k-CITE-seq dataset is presented in Figure 7.a, which shows a good separation between different cell types.

**Figure 6:**
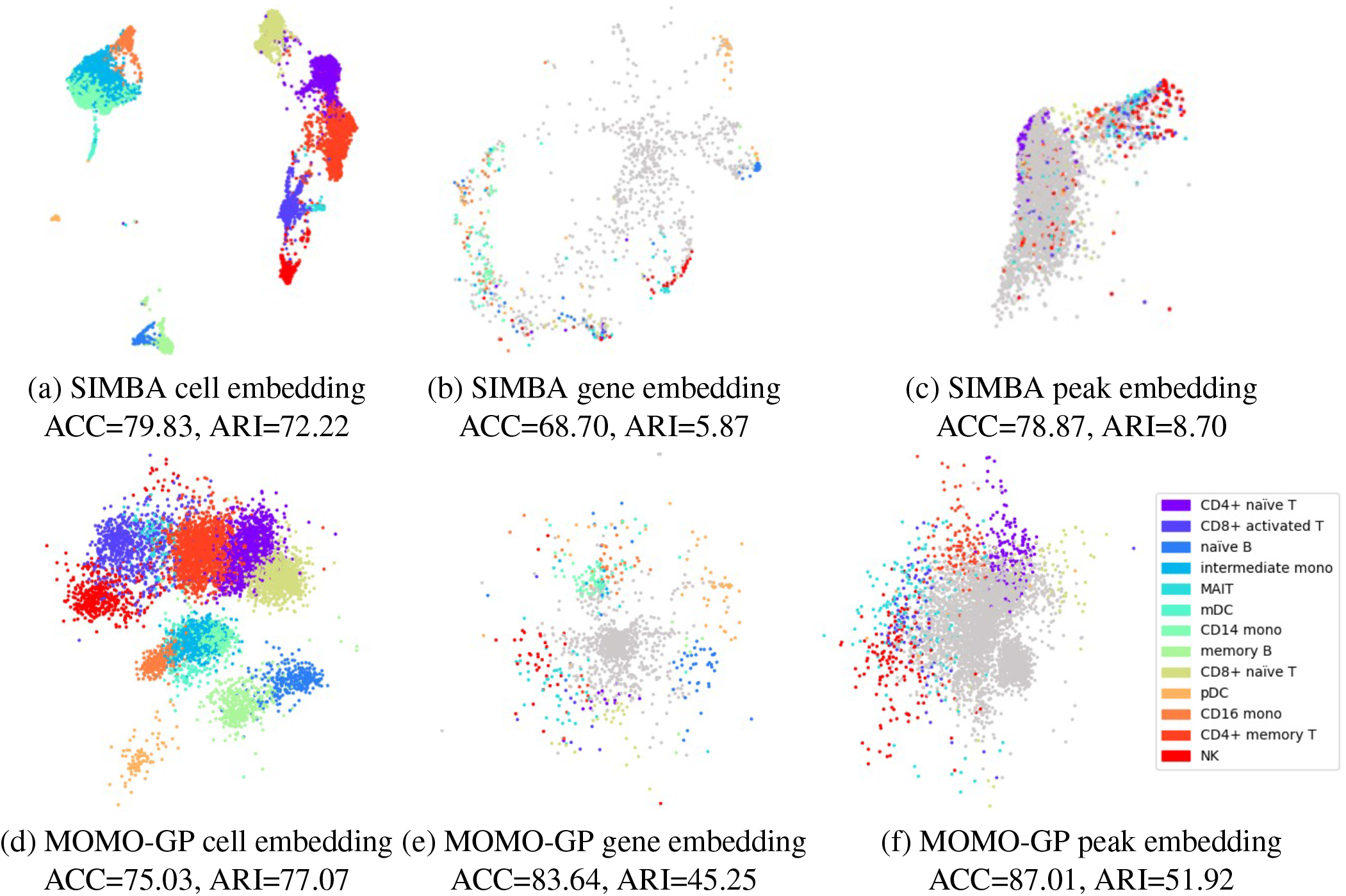
Exploration of the PBMC 10k dataset with SIMBA and MOMO-GP embedding techniques applied to both scRNA-seq and scATAC-seq data: (a) SIMBA-UMAP embedding of cells color-coded by cell types. (b) SIMBA-UMAP embedding of genes, with the top 100 marker genes per cell type colored by their respective cell types. (c) SIMBA-UMAP embedding of peaks, with the top 500 marker peaks per cell type colored by their corresponding cell types. All mappings are conducted in a 50-dimensional space. (d) MOMO-GP embedding of cells, (e) MOMO-GP embedding of genes, and (f) MOMO-GP embedding of peaks, where cells, genes, and peaks are projected into a 2-dimensional space. Non-marker genes and peak are shown in gray. For a more quantitative comparison, ACC and ARI values are also presented.

**Figure 7:**
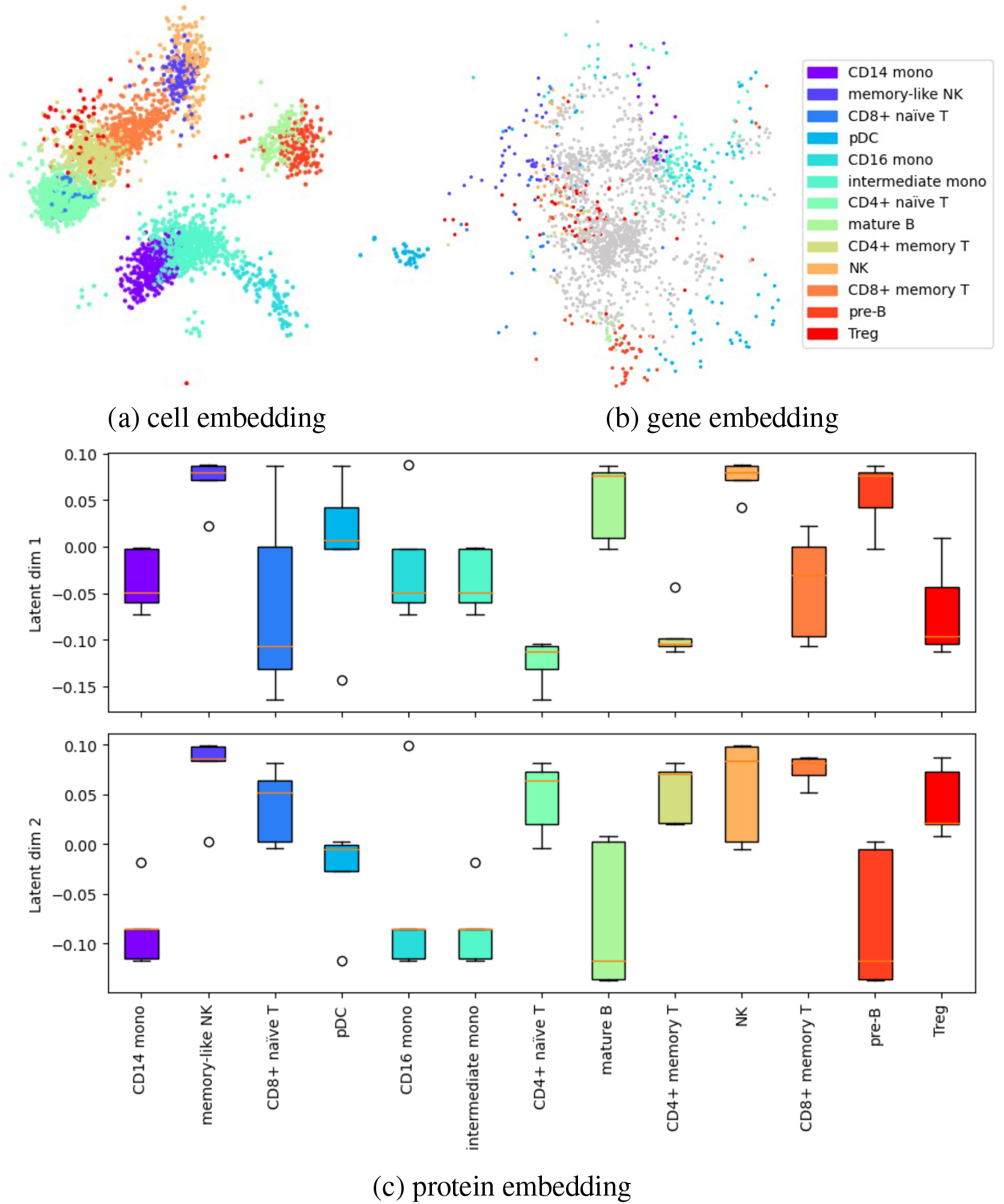
Visualization of MOMO-GP-embedded scRNA-seq data and surface protein data from the PBMC 5k (CITE-seq) dataset, with cells, genes, and proteins mapped to a 2D space: (a) Embedding of cells colored by cell types. (b) Embedding of genes, with the top 100 marker genes in each cell type colored by their corresponding cell type. Non-marker genes are shown in gray. (c) First and second embeddings of proteins, with the top 5 marker proteins considered for each cell type.

In the multi-view version of MOMO-GP, we utilize a single embedding for the samples and apply the MRD approach to assign distinct coordinates to each view, along with some shared coordinates across both views. To illustrate the shared and specific coordinates, we use scRNA-seq and scATAC-seq data from the PBMC 10k dataset, varying the number of latent dimensions from the set {2, 4, 8, 16, 32}. We present the corresponding ARD values, **w**_1_ for scRNA-seq data and **w**_2_ for scATAC-seq data. The results are shown in Figure S7. For each latent dimension, the first bar indicates the ARD values for the scRNA-seq data, while the second bar represents the values for the scATAC-seq data. By analyzing these values and establishing a suitable cutoff, we can identify which dimensions of the cell embedding are specific to the scRNA-seq dataset, which are specific to the scATAC-seq dataset, and which are shared between the two datasets.

In this experiment, when setting the number of latent dimensions to 4, as suggested by the figure, latent variables 3 and 4 are shared between both modalities, latent variable 1 is specific to the scRNA-seq data, and latent variable 2 is absent in both modalities. We provide the cell embedding colored by cell types in Figure S8.a for shared coordinates and Figure S8.b for specific and absent coordinates. The figure demonstrates a clear separation of cell types for coordinates with high ARD values in both modalities. However, for coordinates with low ARD values in either (RNA or ATAC) or both modalities (absent), this separation of cell types is less distinct.

#### 3.2.2 Gene Embedding

In this section, we assess the gene embedding produced by the multi-view MOMO-GP model for both the PBMC 10k and PBMC 5k-CITE-seq datasets.

The results for the PBMC 10k dataset are shown in Figure 6.b and 6.e, while those for the 5k-CITE-seq dataset are presented in Figure 7.b. We conducted a comparison with SIMBA, setting the number of latent dimensions for gene representation to 2 for MOMO-GP and 50 for SIMBA. We visualize the 2D embedding of genes directly obtained from MOMO-GP and the UMAP embedding of SIMBA-embedded data. Embedding of genes into 50-dimensional space using MOMO-GP followed by UMAP is given in Figure S6.b. In these figures, we identify the top 100 differentially expressed marker genes and color them according to their respective cell types. The visualization reveals that our gene embedding, using only 2 latent factors, provides meaningful insights. While there is no perfect separation between marker genes of all cell types, all marker genes of a specific cell type tend to form cohesive clusters (ACC=83.64, ARI=45.25). These results highlight the superior performance of our method over SIMBA’s gene embedding.

#### 3.2.3 Peak Embedding

The peak embeddings generated by both the SIMBA and MOMO-GP models for the PBMC 10k dataset are displayed in Figure 6.c and 6.f, respectively. In these figures, the number of latent dimensions is set to 2 for MOMO-GP and 50 for SIMBA. The results for 50-dimensional space using MOMO-GP is given in Figure S6.c. In these figures, the top 500 marker peaks for each cell type have been selected and colored according to their respective cell types. As observed, our method demonstrates a clear separation between marker peaks, while the results obtained from SIMBA fall short in comparison.

#### 3.2.4 Protein Embedding

The protein embedding generated by our method is depicted in Figure 7.c. Given that the dataset contains a total of 32 proteins, the 2D visualization of data points may not be as informative. Furthermore, due to the overlap between the top 5 marker proteins for different cell types, it was not feasible to color the points based on their relevant cell types. Therefore, we opted to use box plots for this analysis. For each cell type, we selected the top 5 marker proteins and displayed their values for the first and second latent dimensions learned by MOMO-GP. While some overlap exists between relevant cell types such as CD14 mono, CD16 mono, and intermediate mono, there is a clear separation between irrelevant ones, such as B cells and monocyte cells.

#### 3.2.5 Interpretability of the Model

One of the main advantages of our model is its capability to generate embeddings for both samples and features. In the context of multi-view data analysis, such as CITE-seq data, this translates into distinct embeddings for cells, genes, and proteins. As demonstrated in the single-view version, utilizing the gene relevance map allows us to identify clusters of cells and genes with correlated expression patterns.

Given the abundance of genes in CITE-seq data, aggregating genes into meta genes simplifies analysis and facilitates the assessment of their relevance to meta cells. Similarly, leveraging the protein relevance map enables the identification of clusters of cells and proteins with cohesive expression profiles. With 32 different proteins in CITE-seq data, determining the relevance of each protein to a group of cells becomes straightforward. By identifying cells relevant to each protein and cells relevant to each meta gene, we can establish relationships between proteins and genes.

In other types of single-cell omics data, such as scRNA-seq and scATAC-seq, we can similarly uncover relationships between groups of genes and peaks. Overall, the ability of MOMO-GP to generate embeddings for diverse types of data facilitates comprehensive analysis and the discovery of complex relationships between molecular features and cellular phenotypes across various omics datasets.

Figure 8 illustrates the protein relevance map for the PBMC 5k CITE-seq dataset. Among the 32 proteins analyzed, 11 specific proteins have been identified as highly relevant to distinct cell groups, effectively covering all cells in the dataset. For instance, in the first row of the figure, CD16, CD56, and TIGIT exhibit notable relevance in NK and memory-like NK cells. Moving to the second row, CD127, CD28, and CD27 demonstrate significant relevance in memory CD4+ Naïve T, CD8+ Naïve T, memory CD4+ T, and memory CD8+ T cells. In the third row, CD14, CD86, and HLA-DR exhibit the highest relevancy in CD14 monocytes, intermediate monocytes, and CD16 monocytes. Lastly, in the last row, CD19 and CD20 display the highest relevancy in mature B cells and pre-B cells.

**Figure 8:**
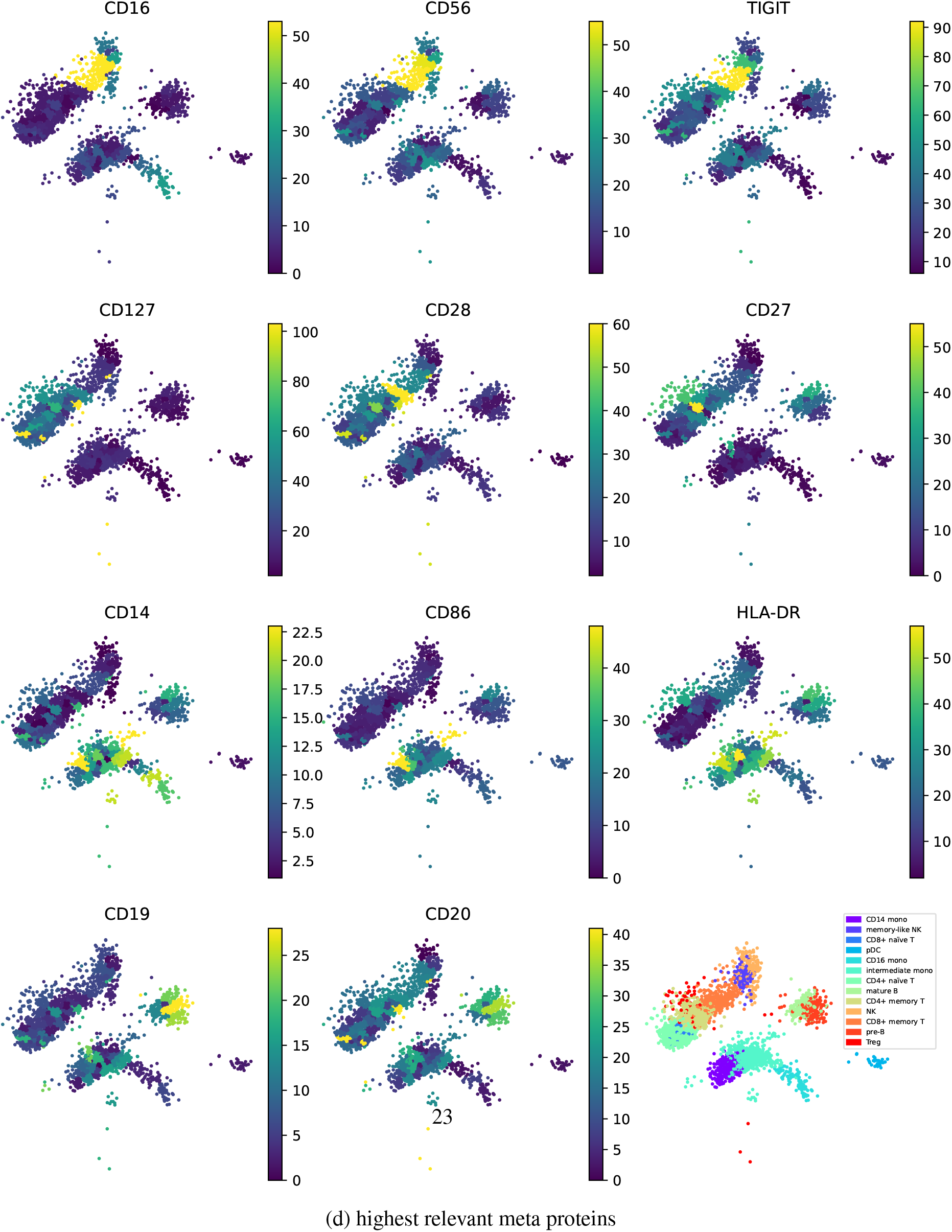
Exploration of the PBMC 5k (CITE-seq) dataset using a protein relevance map, which automatically detects correspondence between groups of cells and proteins. The protein relevance plot highlights areas where the contribution-of proteins is highest. For example, in the first-row, CD 16, CD56, and TIGIT exhibit high relevance in NK and memory-like NK cells. In the second rowrCD127, CD28, and CD27 demonstrate high relevance in memory CD4+ Naive T, CD8+ Naive T, memory CD4+ T, and memory CD8+ T cells. In the third row, CD 14. CD86, and HLA-DR show the highest relevancy in CD 14 monocytes, intermediate monocytes, and CD16 monocytes. In the last row, CD19 and CD20 display the highest relevancy in mature B cells and pre-B cells.

As previously demonstrated in the single-view version, the gene relevance map allows us to determine the relevance of cells and genes. Subsequently, by identifying shared sets of cells relevant to both genes and proteins, we can establish the relevance between genes and proteins. The gene relevance map after running multi-view version of MOMO-GP for the PBMC 5k CITE-seq dataset is shown in Figure S10. In this figure, cells relevant to meta-genes 2 and 3 are indicated by circles. Additionally, the protein relevance map in Figure 8 shows that proteins CD16, CD56, and TIGIT are relevant to these cells. This suggests a potential relevance between meta-genes 2 and 3 and these proteins. This claim is validated as these proteins are top markers for NK and memory-like NK cells, and the genes related to meta-genes 2 and 3 are marker genes for cells with these NK and memory-like NK cell types.

## 4 DISCUSSION

In this section, we discuss the properties of MOMO-GP and potential areas for improvement. MOMO-GP is a novel multi-view latent variable model that captures the nonlinear structure of data by combining a flexible embedding layer with a Gaussian Process layer. It learns separate latent representations for cells and features (such as genes, peaks, proteins, etc.) in an interpretable manner. By embedding features and utilizing the concept of a gene relevance map, we can identify groups of cells and correlated features from different modalities.

The closest work to ours, which learns sample and feature embeddings, is SIMBA [10]. We compare our results with SIMBA in both single-view and multi-view settings, demonstrating that the feature embeddings learned by our method are more meaningful. In contrast with SIMBA, which uses a single embedding for both samples and features, our method employs different embeddings for each. This enhances the interpretability of the model, as the gene embeddings in MOMO-GP link more naturally to the cell embeddings and perform better than those in SIMBA. We demonstrate this outperformance using various visualization plots and by providing accuracy (ACC) and Adjusted Rand Index (ARI) values.

Proposing a Bayesian version of the model and placing a prior on the latent variables could be explored in future research. Another direction for future work is to place a neural network on top of the embeddings to better capture the non-linear structure of the data. Considering sample-based data such as time series data or spatial transcriptomic data, where proximity information is crucial, could be another direction for future research. To address this, we should further develop the neural network layer to effectively handle such datasets.

## 5 CONCLUSION

In this paper, we introduced a new method called Multi-Omics Multi-Output Gaussian Processes (MOMO-GP) for integrating multi-omics data. The key feature of this method is its ability to simultaneously learn separate feature embeddings for each modality and a shared sample embedding. This approach strikes a balance between expressive power and interpretability. The expressive power is enhanced by linking the embedding layer to the Gaussian Process layer, while interpretability is achieved by explicitly modeling non-linear dependencies between features and samples. Through various experiments, we demonstrated that our model outperforms existing algorithms.

## 6 DATA AVAILABILITY

The code for MOMO-GP is available at https://github.com/MLO-lab/MOMO-GP

## 7 ACKNOWLEDGEMENTS

We extend our gratitude to the researchers of the Machine Learning in Oncology (MLO) Lab at DKTK/DKFZ and Frankfurt University Hospital for their invaluable contributions during numerous talks and discussions on this project.

Co-funded by the European Union (ERC, TAIPO, 101088594). Views and opinions expressed are however those of the authors only and do not necessarily reflect those of the European Union or the European Research Council. Neither the European Union nor the granting authority can be held responsible for them.

## 7.0.1 Conflict of interest statement

FB reports funding from Merck KGaA and Bayer AG and renumeration from Albireo and Siemens AG.

## 8 SUPPLEMENTARY METHODS

### 8.1 Mathematical Foundations of the Single-View Probabilistic Modeling of MOMO-GP

In this section, we elaborate on the probability distribution and the dependencies between different variables of single-view MOMO-GP. Assuming **Y** *∈* ℝ^*I×J*^ represents the given dataset, where *I* is the number of data points and *J* is the number of attributes. In real-world applications, it is common to consider the observed data points **Y** as noisy measurements of true values **F**. This assumption leads to a factorized likelihood given by:

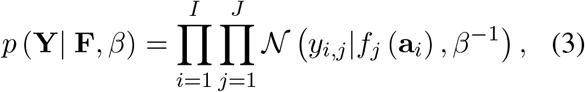

where *f*_*j*_ (**a**_*i*_) represents a function of an *r*-dimensional latent space. The points in the lower-dimensional latent space are represented by the matrix 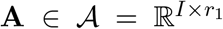. The relationship between the latent space and the data space is defined by the function *f*_*j*_ (**a**_*i*_).

We choose a Gaussian process prior for **F** *∈* ℝ^*I×J*^. Using the idea of GP-LVM, we fit *J* independent GP regression models on the unobserved latent variable **a**_*i*_, with

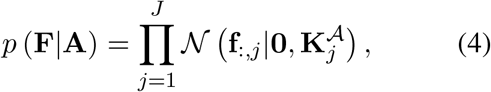

where 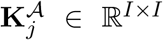 defines the covariance between each pair of **A**, i.e., (**a**_*i*_, **a**_*i*_*′*). Here, **F** is given as the output, and the latent representation **A** is optimized.

Now, we deviate from the assumption of independence among *J* different attributes, recognizing that this assumption is not always accurate. To address this, we introduce a new coregionalization kernel for performing multi-output Gaussian Process regression modeling. This kernel is a separable kernel expressed as the Kronecker product of two individual kernels. The first kernel measures the similarity of samples in the input space, while the second kernel captures the similarity between each pair of features in lower-dimensional space 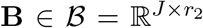, i.e. (**b**_*j*_, **b**_*j*_*′*).

The first kernel, denoted as **K**^*𝒜*^ *∈* ℝ^*I×I*^, defines the covariance between each pair of **A**. Similarly, the second kernel, denoted as **K**^*ℬ*^*∈* ℝ^*J ×J*^, defines the covariance between each pair of **B**. This concept is derived from LVMOGP, where in their supervised scenario, inputs are observed. However, in our model, both kernels act on latent variables **A** and **B**, and they need to be optimized.

The proposed kernel can be expressed as:

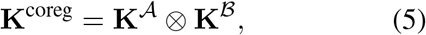

where

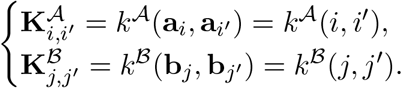

In this manner, we consider a correlation between two entities (*i, j*) and (*i*^*′*^, *j*^*′*^) in matrix **F**. This correlation is defined by two kernels over the latent spaces *α* and *β* as follows:

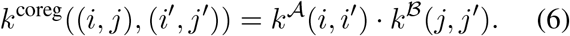

The size of this coregionalization kernel is (*I* · *J*) *×* (*I* · *J*) since it computes the correlation for all combinations of *I* samples and *J* features in matrix **F**. Finally, by concatenating all entities of matrix **Y**, we can model our observed data (*i, j, y*_*i,j*_) with a Gaussian Process as follows:

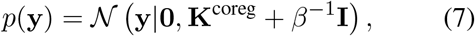

where **y** *∈* ℝ^*I*·*J*^ is the noisy version of **f** *∈* ℝ^*I*·*J*^ and **K**^coreg^ *∈* ℝ ^(*I*·*J*)*×*(*I*·*J*)^.

To compute this GP model, we need to compute the inverse of the covariance matrix **K**^coreg^, which has a complexity of 𝒪 (*n*^3^), where *n* is the number of samples, in this case *n* = *I* · *J*. In our application, such as gene expression data, we often have a large number of cells (*I*) and genes (*J*). To decrease the time complexity of the model and make the problem tractable, we need to employ the idea of sparse GPs.

In sparse GPs, the concept is to expand the probability space with *n*^*′*^ different auxiliary pairs of input-output variables collected in matrices 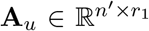 and **u *∈*** ℝ^*n′×J*^. Here, the inducing output variables **u** are assumed to have the same GP prior as the variables **f**. The prior over these variables therefore takes the form:

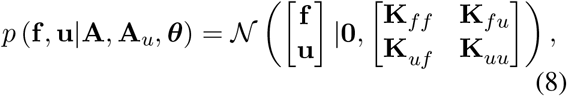

where **K**_*ff*_ is built by computing the covariance function on all latent variables **A, K**_*uu*_ is constructed by evaluating the covariance function on all auxiliary samples **A**_*u*_, **K**_*fu*_ is the cross-covariance between latent variables and auxiliary samples, and 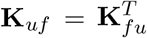. The dependence on variables **A, A**_*u*_, and the parameters ***θ*** is through these kernel matrices.

The Gaussian process definition allows us to write the marginal distribution and conditional distributions as follows [13]:

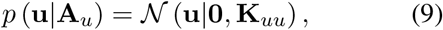

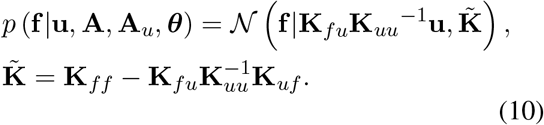

By utilizing equations (9) and (10), the marginal distribution can be expressed as:

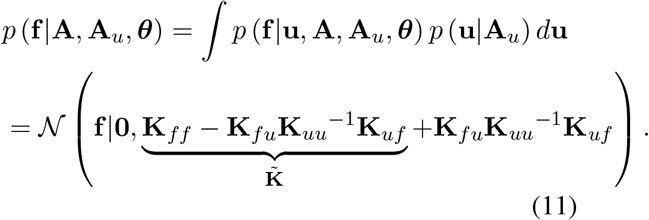

Using an approximate posterior, we have:

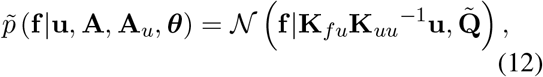

where 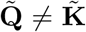, and thus the marginal distribution would be:

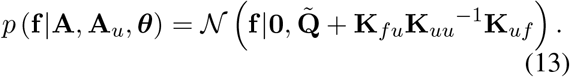

There are different kinds of approximation for 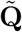. For example, in conditional (DTC) approximation 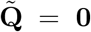 [38, 39]. In our implementation, we use the Scalable Variational Gaussian Process (SVGP) model [40] to speed up the training. These sparse GP models are associated with a computational cost of *O (nn*^*′*2^), *n*^*′*^ ≪ *n* [40]. Decreasing the time complexity is because of avoiding computation and inversion of full covariance matrix **K**_*ff*_, and instead calculation of matrices **K**_*fu*_ and **K**_*uu*_.

In our coregionalization kernel,

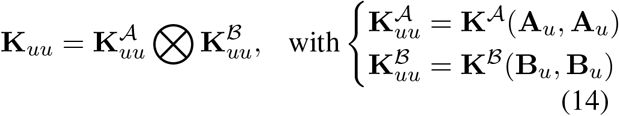

and

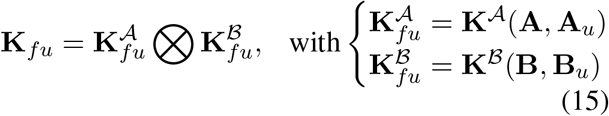

In this model, we define *m*_*A*_ inducing points **A**_*u*_ for the sample space and *m*_*B*_ inducing points **B**_*u*_ for the feature space. The matrices 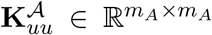 (resp. 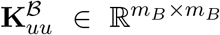) compute the similarity between all inducing variables **A**_*u*_ (resp. **B**_*u*_) in space *𝒜* (resp. in space *ℬ*), and the matrices 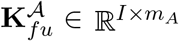 (resp. 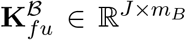) compute the cross-covariance between latent variables **A** and **A**_*u*_ (resp. **B** and **B**_*u*_).

By leveraging the concept of sparse GP, we reduce the time complexity of our model from 𝒪 ((*I* ·*J*))^3^ to 𝒪 ((*I* ·*J*) (*m*_*A*_ ·*m*_*B*_)^2^). Moreover, we enforce the same number of inducing points for **A**_*u*_ and **B**_*u*_ and that allows us to replace the Kronecker product with an elementwise product [18]. Using this, we can then further reduce computational complexity to 𝒪 ((*I* ·*J*) *m*^2^)).

In our model, the variables that need to be optimized include **A, A**_*u*_, **B, B**_*u*_, and other kernel parameters. A novel aspect of our model is the combination of an embedding layer with a Gaussian Process layer to capture the nonlinear structure of the data. Instead of directly optimizing variables **A** and **B**, we utilize an embedding function that converts positive integers (indexes) into dense vectors of fixed size.

To learn **A** and **B**, we map all indices in the range 1, …, *I* and 1, …, *J* to matrices of size *I × r*_1_ and *J × r*_2_, respectively, using an embedding layer. Here, *r*_1_ represents the size of the input embedding space, and *r*_2_ represents the size of the output embedding space. For computing **A**_*u*_ and **B**_*u*_, we randomly selec, t from 1, …, *I* and 1, …, *J*, respectively, and pass them through the embedding layer to obtain matrices **A**_*u*_ and **B**_*u*_ of size *m*_*A*_ × *r*_1_ and *m*_*B*_ × *r*_2_, respectively. During training, the weights of this embedding layer are optimized.

#### 8.1.1 Coupling inducing points

In traditional multi-output GP formulations, the covariance matrix of the inducing variables, **K**_*uu*_, is computed as 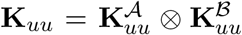. By selecting the same number of inducing inputs for **A**_*u*_ and **B**_*u*_, we can couple the inducing points for both inputs and outputs, allowing us to reformulate the construction of **K**_*uu*_, **K**_*uf*_, and **K**_*fu*_. Specifically, when computing the cross-covariance between the *q*-th training sample (*i, j*) and the *l*-th pair of inducing points to build **K**_*fu*_, we obtain:

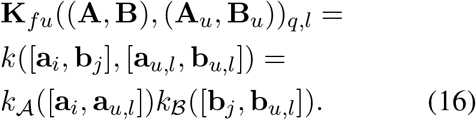

This means that by using paired inducing points, **K**_*fu*_ can be written as the elementwise product between 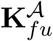 and 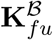.

We construct the covariance matrix **K**_*uu*_ using the same pairing approach, computing the covariance between the *o*-th and *p*-th inducing points as:

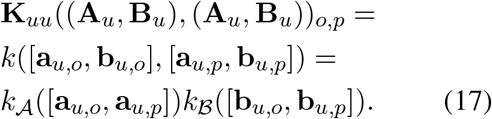

So, **K**_*uu*_ is writen as elementwise product between 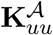 and 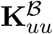. By these formulations, the time complexity of sparse multi-output GP will decrease to 𝒪 ((*I* ·*J*) *m*^2^). Algorithm 1 summarizes all the steps involved in the implementation.

##### Algorithm 1

MOMO-GP (Multi-Omics Multi-Output Gaussian Process algorithm for embedding both samples and features)

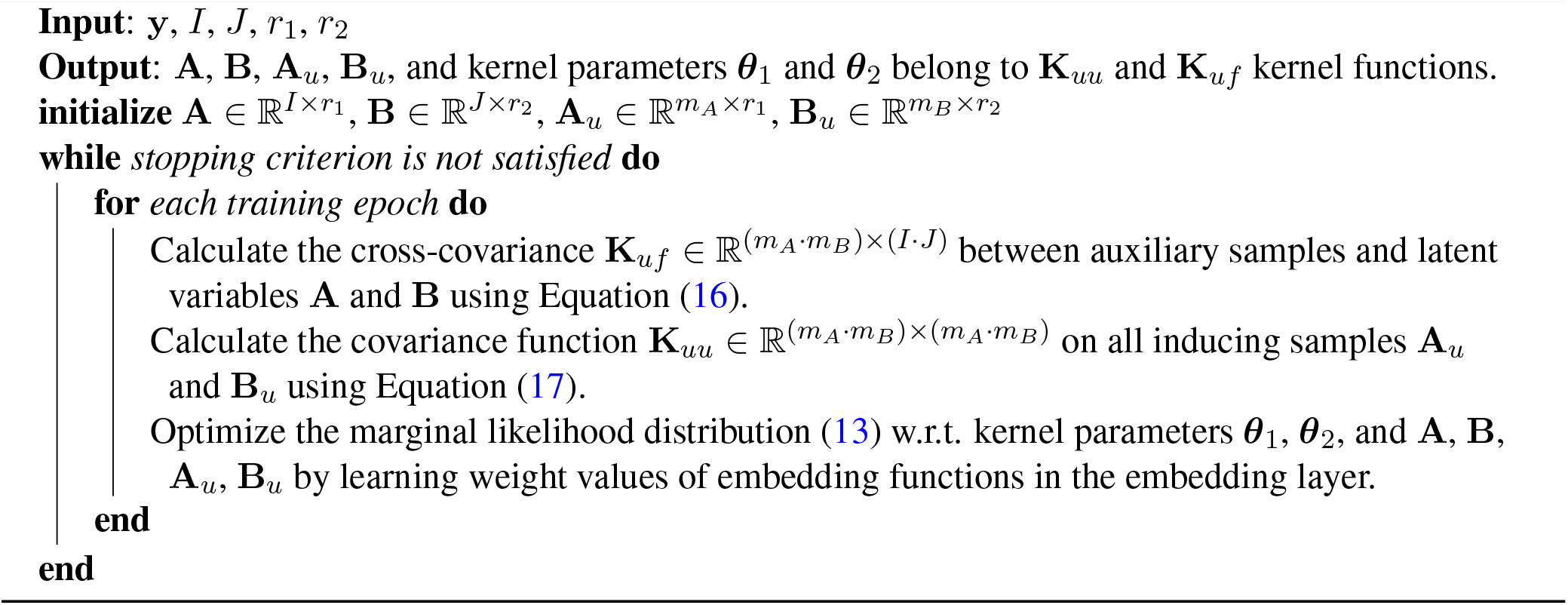

### 8.2 Gene relevance maps

The differential *d*_*gc*_ for each gene *g* and cell *c* describes the change in gene expression space *Y* along the embedding space *A*. The partial derivative of *d*_*gc*_ for each dimension *r* ***∈*** *{*1, …, *R}* is defined as follows:

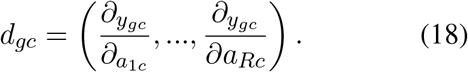

When computing this derivation is not mathematically possible, we can use the estimation 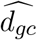 for each cell *c* using its *k* nearest neighbors *n* ∈ NN_*k*_(*c*) as follows:

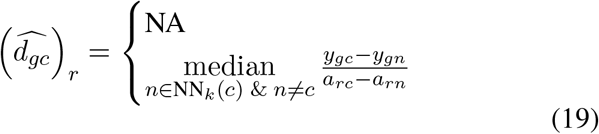

The Euclidean norm of *d*_*gc*_ for all dimensions *r* can then be defined as follows:

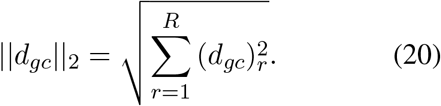

For each cell *c*, all the genes should be ranked according to the values of ||*d*_*gc*_||_2_ from high to low. If we select a collection of cells Ψ *⊆ {*1, …, *C}*, define a rank cutoff rg_*max*_, and have the rank of genes rg according to the values ||*d*_*gc*_ ||_2_, then we can define the local gene relevance values for each subset Ψ as follows:

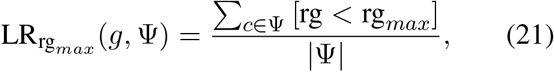

in which bracket notation

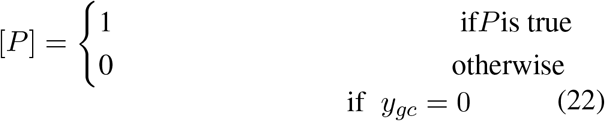

The local gene relevance value LR(*g*, Ψ) shows if the contribution of gene *g* in area Ψ is high or not. Global gene relevance 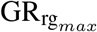 is simply defined by LR (*g*, Ψ) if Ψ is the set of all cells:

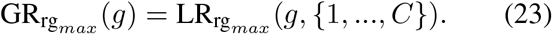

Finally, a gene relevance map identifies the areas in which the contribution of a subset of genes is highest. These genes can be selected from those with the highest global relevance values.

## 9 Data pre-processing

For preprocessing the two datasets, we used Scanpy [24] to perform normalization, logarithmic transformation, clustering, and cluster annotations. The specific preprocessing steps for each dataset are described in detail below:

### Single-cell RNA-seq

We applied quality control measures to the RNA data, filtering out cells and genes that did not meet predefined quality thresholds based on standard criteria. We filtered cells based on the number of detected genes: cells with fewer genes than a set threshold 200 were considered low quality, while those with more than the threshold 5000 were likely doublets. Next, we filtered cells based on the total RNA molecule count (UMI), keeping only cells with fewer than 15,000 counts. Then, we filter cells with high percentages of mitochondrial gene expression by keeping only cells with less than 20 percent mitochondrial content. Additionally, we excluded genes detected in fewer than a certain number of cells (less than 3 cells), focusing the analysis on more widely expressed and biologically relevant genes. Subsequently, the data underwent normalization and logarithmic transformation. To annotate cell types, we employed Leiden clustering, and based on the top marker genes of each cluster, we assigned the corresponding cell type. Clusters displaying noise or exhibiting elevated ribosomal gene expression compared to others, or comprising proliferating cells, were excluded. After these procedures, we performed an additional feature selection step to filter genes based on their Coefficient of Variation (CV) relative to their mean expression. This further reduced the number of genes included in our analysis to 2000, ensuring that only the most variable and informative genes were retained for downstream analysis.

### Single-cell ATAC-seq

For the ATAC-seq data, we initially filtered out cells with insufficient peaks and those with peaks detected in too few cells. Specifically, we retained only the peaks present in at least 10 cells for further analysis. Next, we filtered cells based on the number of accessible chromatin regions (peaks) detected per cell, keeping only those with peak counts exceeding 2000 and less than 15,000. Finally, we excluded cells with total counts below 4000 or above 40,000.Regarding normalization, we initially applied the Latent Semantic Indexing (LSI) [41] method, commonly employed in processing ATAC-seq datasets. Subsequently, we applied the same log-normalization procedure utilized in scRNA-seq analysis. Cell type annotation was performed using Leiden clustering, where clusters were annotated based on their marker genes, with some clusters being removed and others annotated accordingly. After these procedures, we conducted an additional feature selection step to filter peaks based on their Coefficient of Variation (CV) in relation to their mean expression. This process further reduced the number of peaks included in our analysis to 5000. Only cells passing the respective quality control criteria were retained in each modality. For integration purposes, only cells present in both modalities were considered.

### Single-cell Protein Data

For the protein expression data, we employed the Denoised and Scaled by Background (DSB) method to normalize and denoise the data from droplet-based single-cell experiments [25]. The dataset comprises 32 proteins, and the number of cells in the intersection of the RNA-seq and protein expression data, utilized in our analysis of the 5k PBMCs CITE-seq dataset, amounted to 3891.

## 10 Evaluation metrics

### 10.1 Rand Index and Adjusted Rand Index

The Rand Index computes the similarity between two different clusterings. We should count all pairs of data points which are from the same cluster or different clusters in two different clusterings. Given a set *S* of *n* different elements and two partitions of *S, X* = {*X*_1_, …, *X*_*r*_ *}*, a partition of *S* into *r* subsets, and *Y* = {*Y*_1_, …, *Y*_*s*_ *}*, a partition of *S* into *s* subsets. Then, we define:

- *a* as the number of pairs of elements in *S* which are in the same cluster in *X* and in the same cluster in *Y*,
- *b* as the number of pairs of elements which are in different clusters in *X* and in different clusters in *Y*,
- *c* as the number of pairs of elements which are in the same cluster in *X* and in different clusters in *Y*,
- *d* as the number of pairs of elements which are in different clusters in *X* and in the same cluster in *Y*.

Using these notations, the Rand Index is calculated as follows:

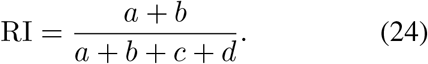

The values of the Rand Index range from 0 to 1, where a value of 0 indicates that the output of both clusterings is completely different, and a value of 1 indicates that both clusterings create exactly the same result.

The Adjusted Rand Index is an extension of the Rand Index. It ranges from -1 to 1, where a value of 1 indicates identical clusterings, a value of 0 indicates random clusterings, and a value of -1 indicates complete disagreement between clusterings. The ARI formula is as follows:

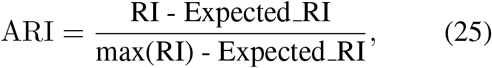

where Expected RI is the expected value of the Rand Index using the permutation model for clusterings.

## 11 Time Complexity of the Model

The time complexity of our model increases linearly with the number of entities in the observed data. To evaluate this, we tested MOMO-GP on single-cell RNA sequencing (scRNA-seq) data from a 5k PBMC CITE-seq dataset. We sampled between 400 and 4000 cells and between 100 and 2000 genes. We then tested all combinations of these cell and gene sets. For each test, we fixed the iteration number at 200, reduced dimensionality to 2, and set the epoch size to 10,000. In this setting, 200 iterations are sufficient for the convergence of the largest dataset. The results are shown in Figure S11. As demonstrated, increasing the size of the observed data results in a linear increase in the model’s time complexity.

## 12 SUPPLEMENTARY FIGURES

**Figure S1:**
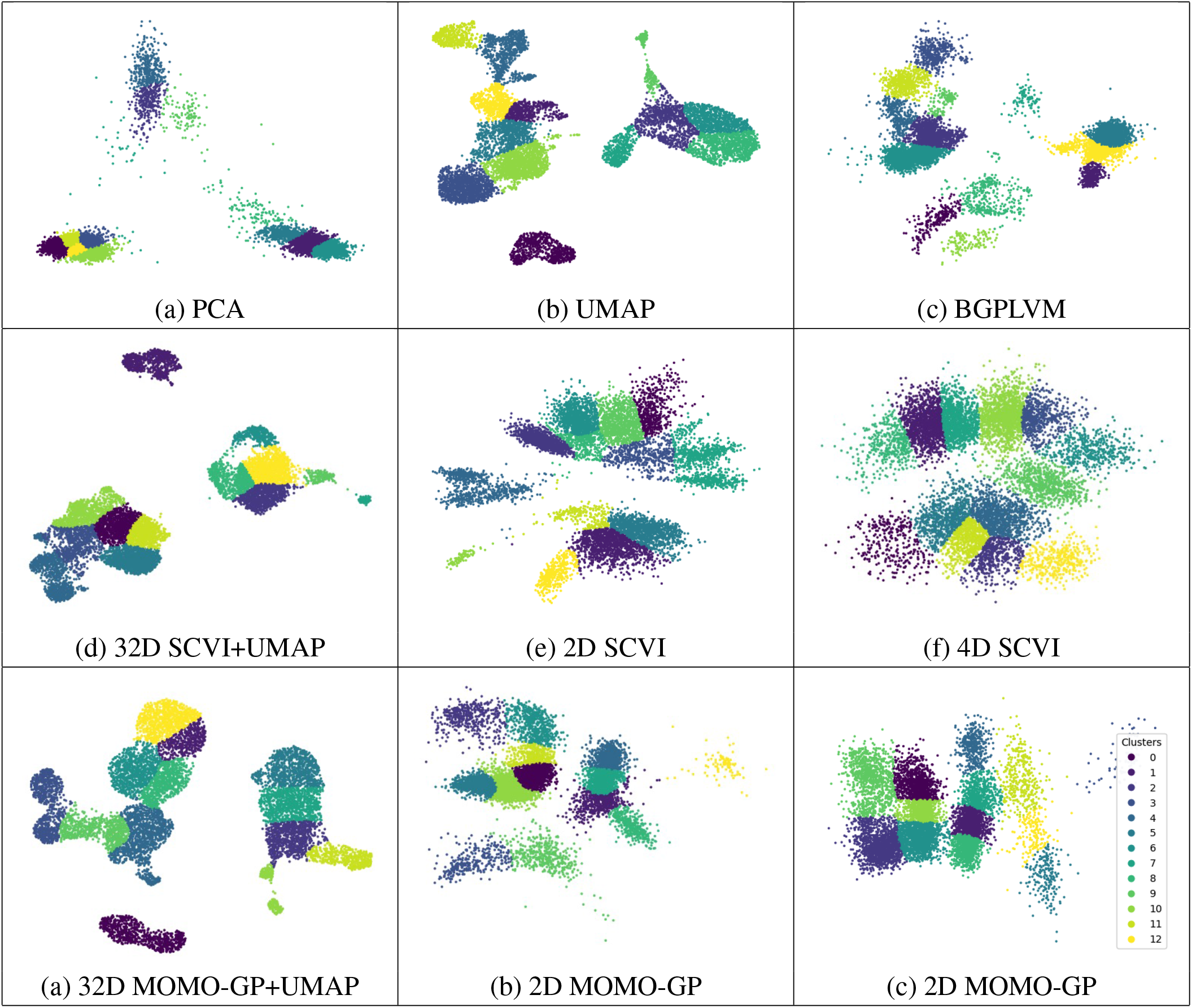
2D visualization of cells colored by clusters in the PBMC 10k dataset using various methods: (a) 2D PCA, (b) 2D UMAP, (c) 2D BGPLVM, (d) 32D SCVI+UMAP, (e) 2D SCVI, (f) 4D SCVI, (g) 32D MOMO-GP+UMAP, (h) 2D MOMO-GP, and (i) 4D MOMO-GP.

**Figure S2:**
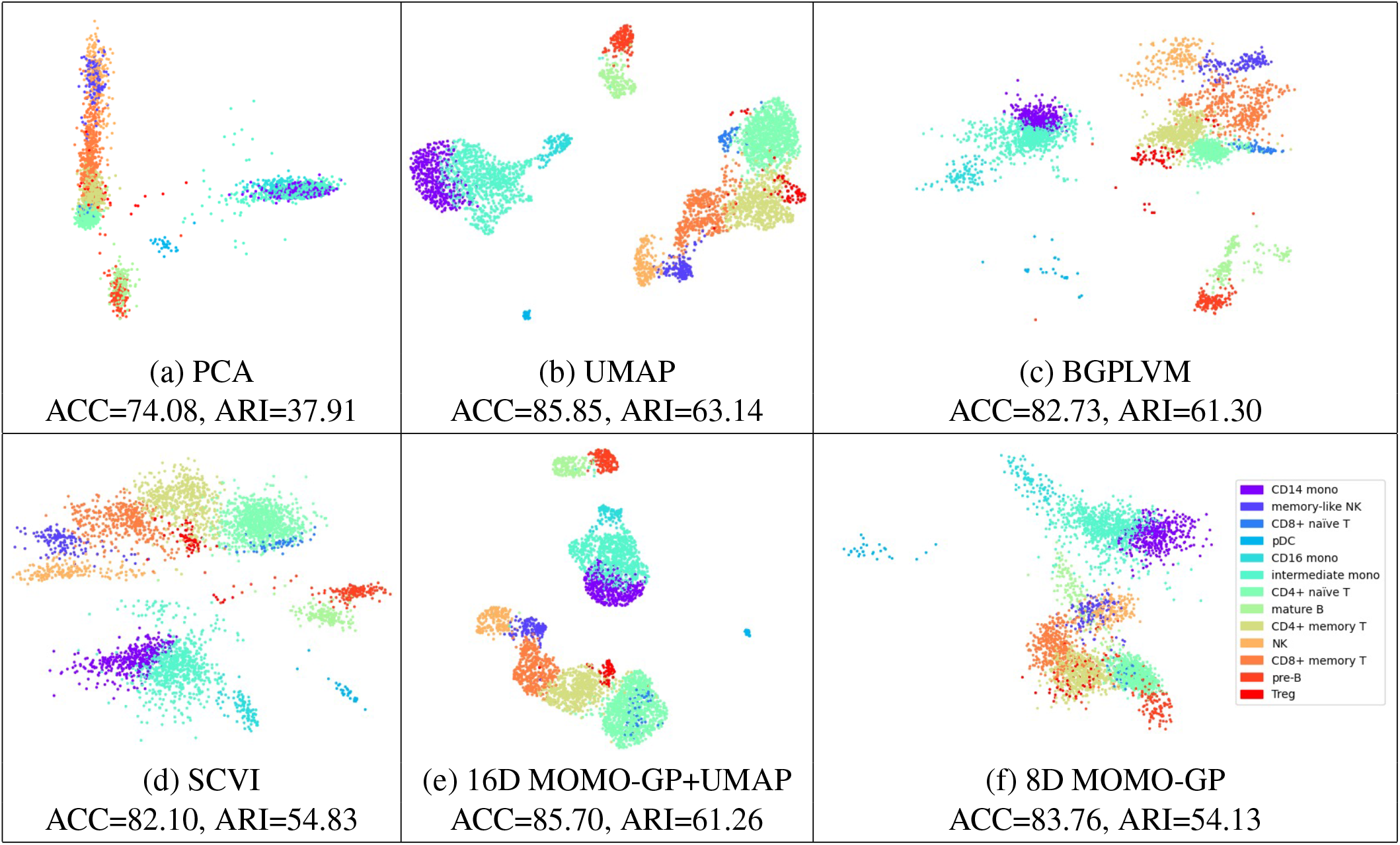
2D visualization of cells in PBMC 5k (CITE-seq) dataset for scRNA-seq data using various methods: (a) PCA, (b) UMAP, (c) BGPLVM, (d) SCVI, (e) 16D MOMO-GP+UMAP, and (f) 8D MOMO-GP+UMAP.

**Figure S3:**
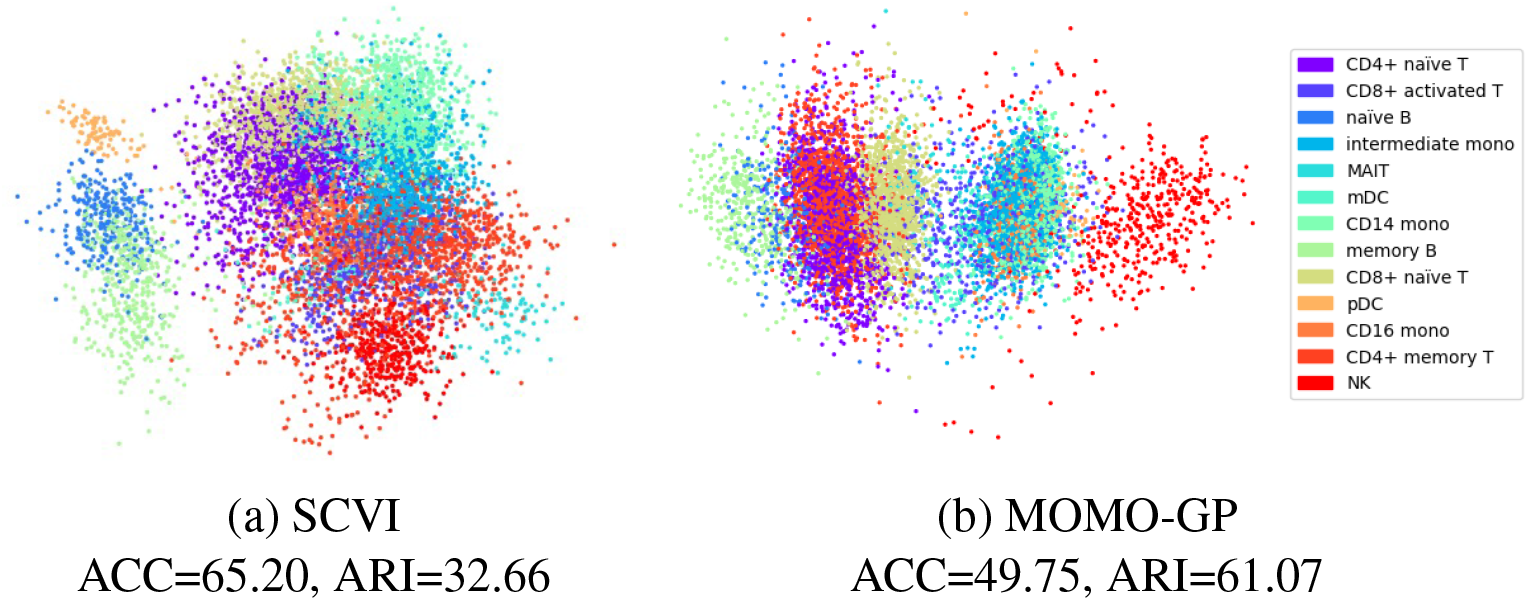
2D visualization of cells in the PBMC 10k dataset for scRNA-seq data using: a) 4D SCVI, and b) 4D MOMO-GP.

**Figure S4:**
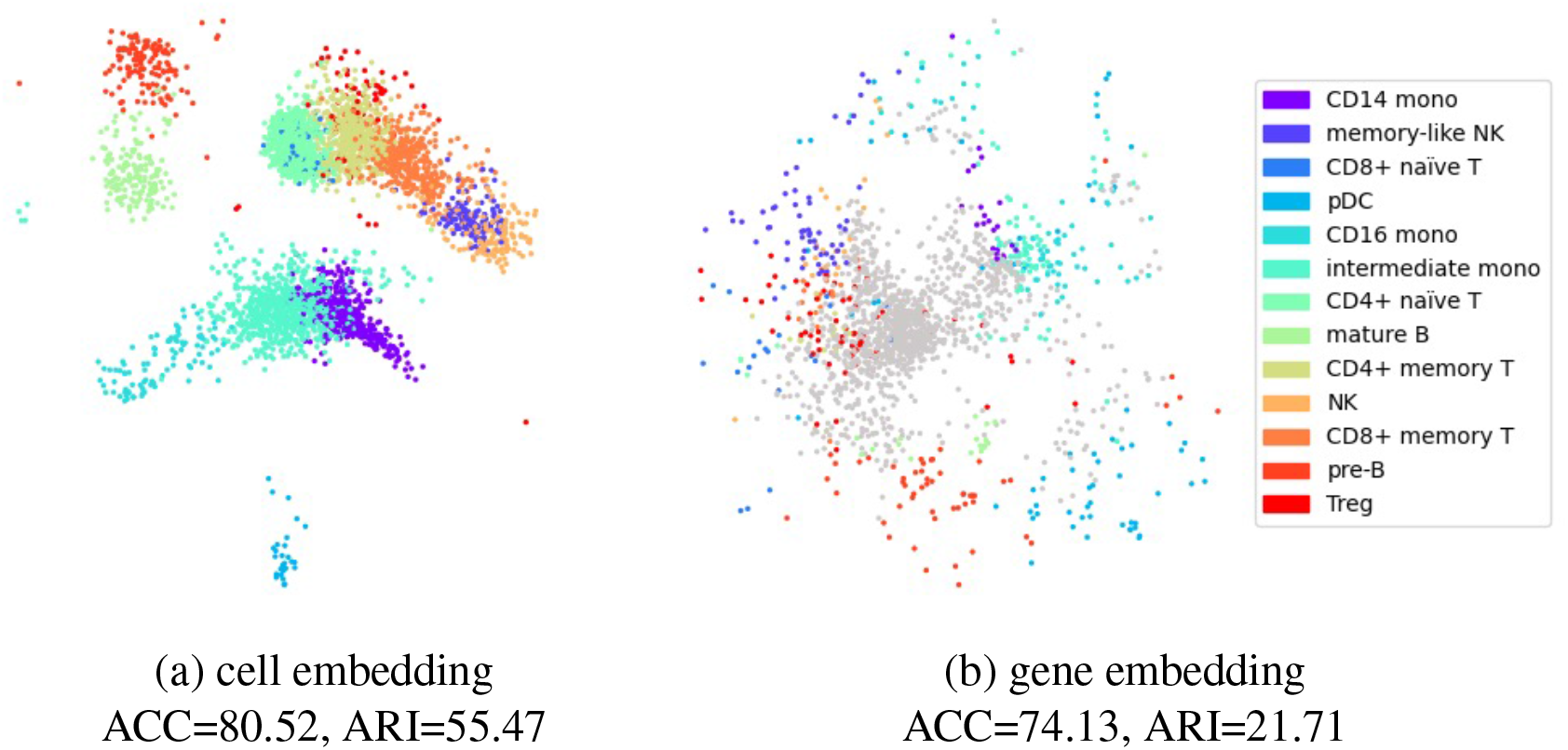
Visualization of MOMO-GP-embedded scRNA-seq data from the PBMC 5k (CITE-seq) dataset, where both cells and genes are mapped to a 2D space: (a) Embedding of cells colored by cell types, (b) Embedding of genes with the top 100 marker genes in each cell type colored by their corresponding cell type. Non-marker genes are shown in gray.

**Figure S5:**
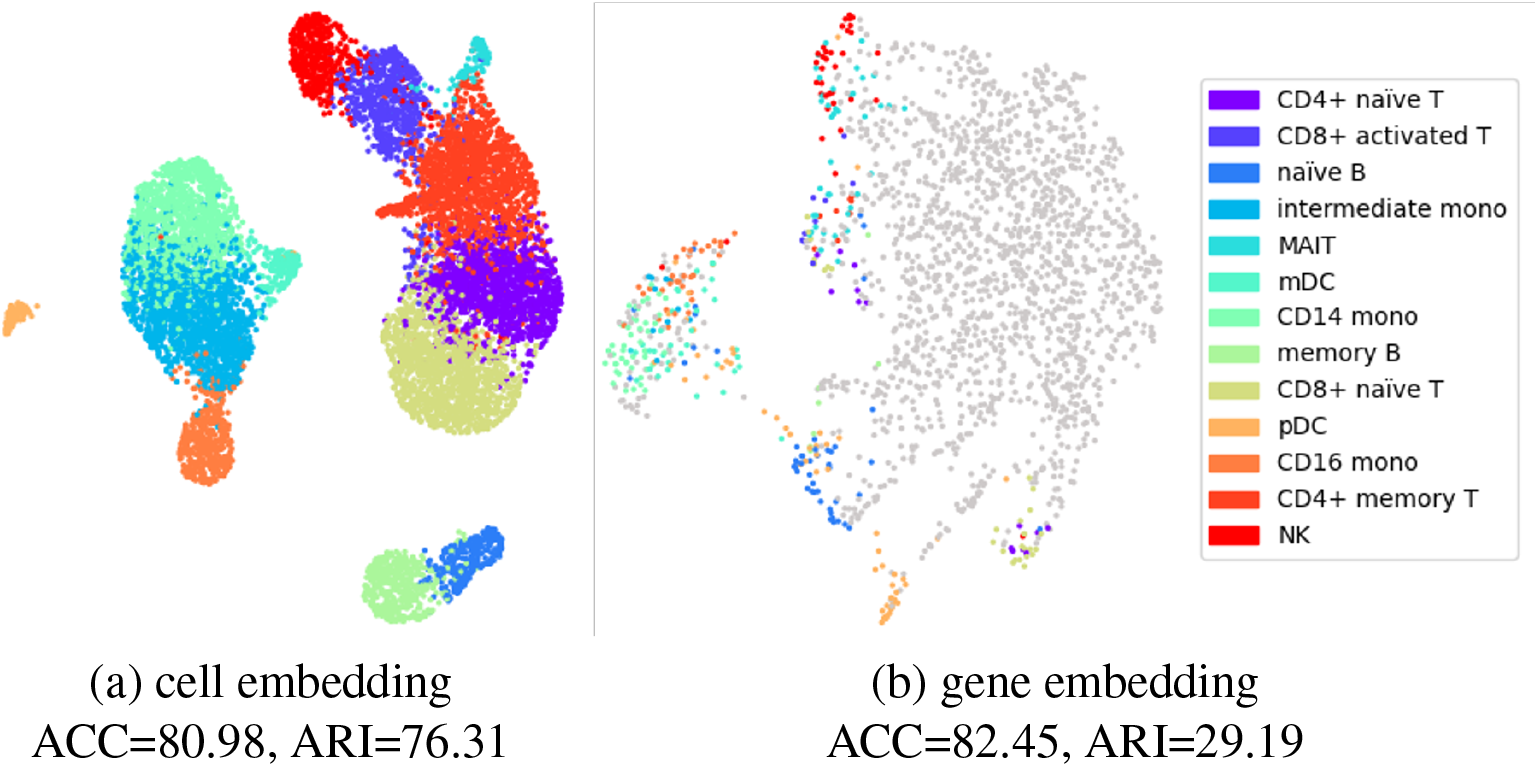
Visualization of PBMC 10k dataset using MOMO-GP embedding techniques for scRNA-seq data. (a) MOMO-GP-UMAP embedding of cells, with cell types color-coded, in a 50-D space. (b) MOMO-GP-UMAP embedding of genes, highlighting the top 100 marker genes per cell type, color-coded by their respective cell types, in a 50-D space. Non-marker genes are shown in gray.

**Figure S6:**
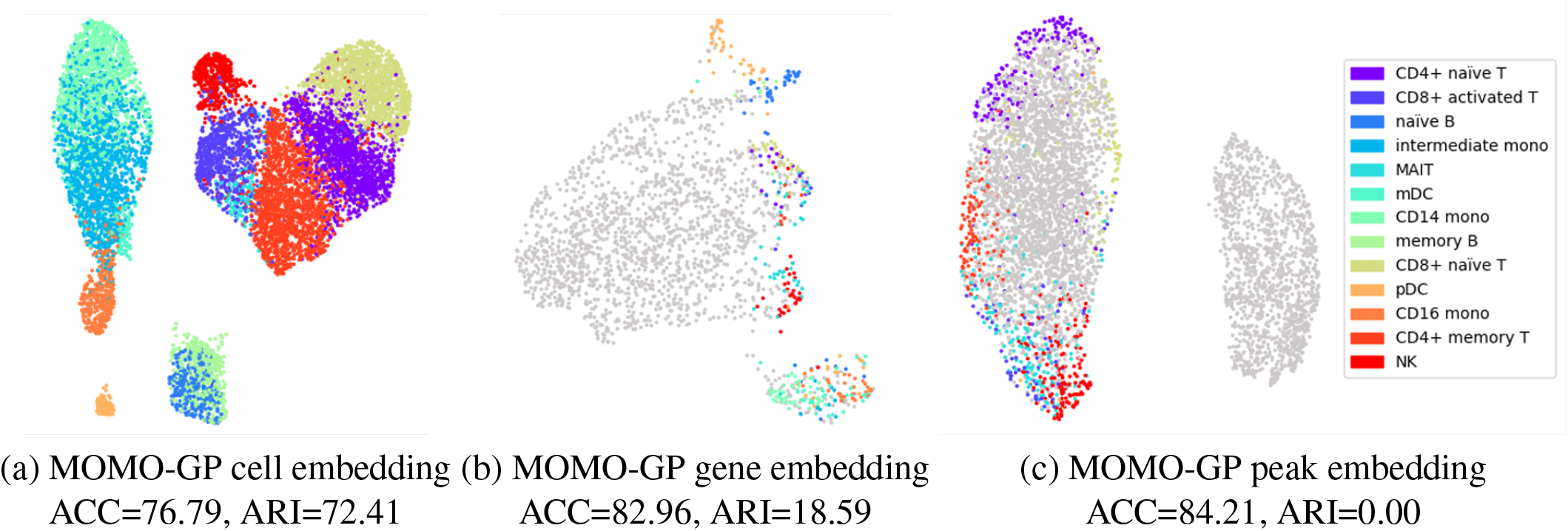
Exploration of the PBMC 10k dataset with MOMO-GP embedding techniques applied to both scRNA-seq and scATAC-seq data: (a) MOMO-GP embedding of cells, (b) MOMO-GP embedding of genes, and (c) MOMO-GP embedding of peaks, where cells, genes, and peaks are projected into a 50-dimensional space using MOMO-GP followed by UMAP. Non-marker genes and peak are shown in gray.

**Figure S7:**
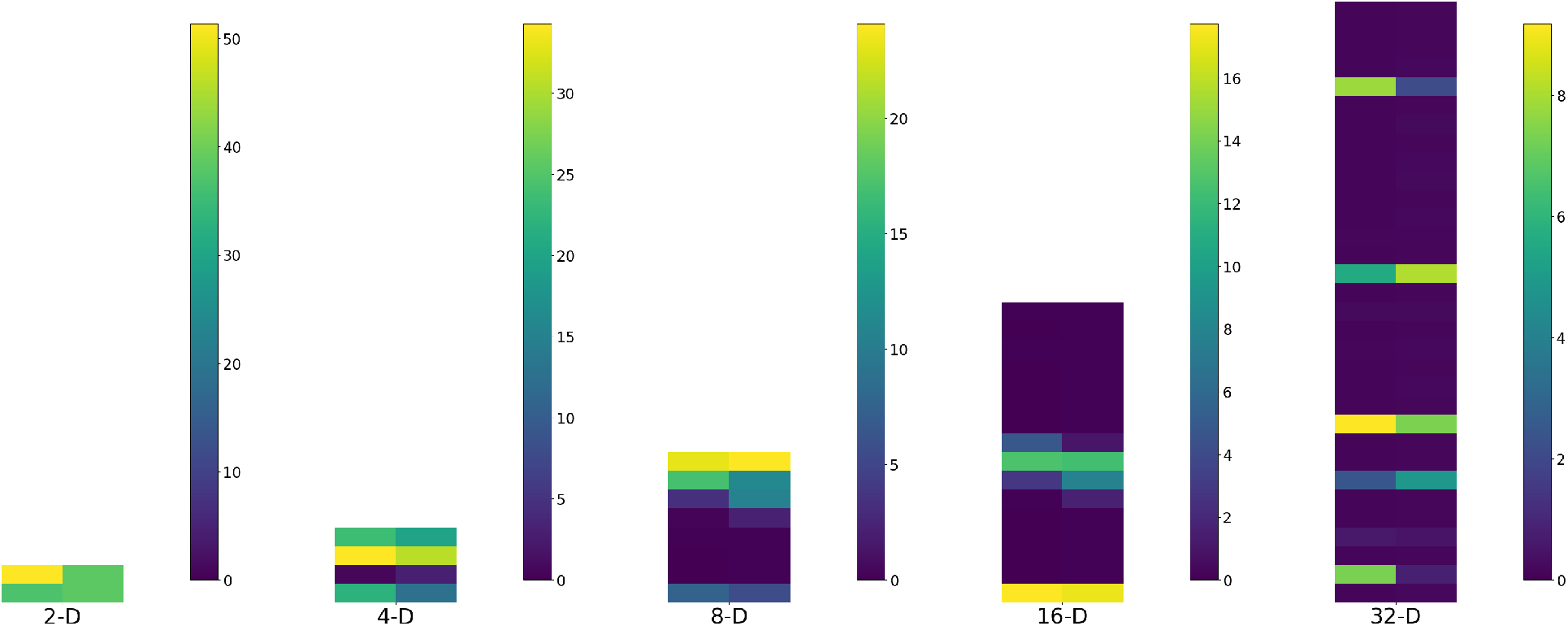
The corresponding ARD values, **w**_1_ for the scRNA-seq data and **w**_2_ for the scATAC-seq data from PBMC 10k dataset. We change the values of different latent dimensions from the set {2, 4, 8, 16, 32 }. For each latent dimension, the first bar represents the ARD values for the scRNA-seq data, while the second bar represents the values for the scATAC-seq data. By comparing these values, we can determine which dimensions of the cell embedding are specific to the scRNA-seq dataset, which are specific to the scATAC-seq dataset, and which are shared between both datasets.

**Figure S8:**
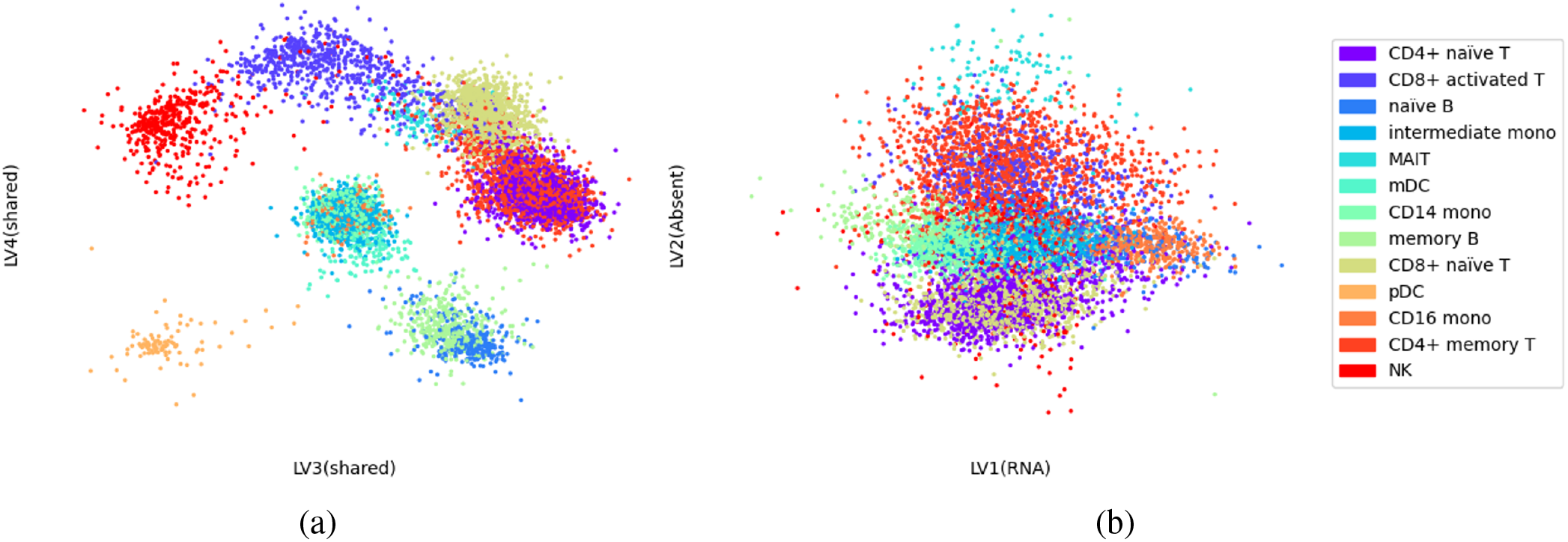
MOMO-GP embedding of cells from the PBMC 10k dataset applied to both scRNA-seq and scATAC-seq data: (a) cell embedding with shared latent variables, (b) cell embedding with specific and absent latent variables.

**Figure S9:**
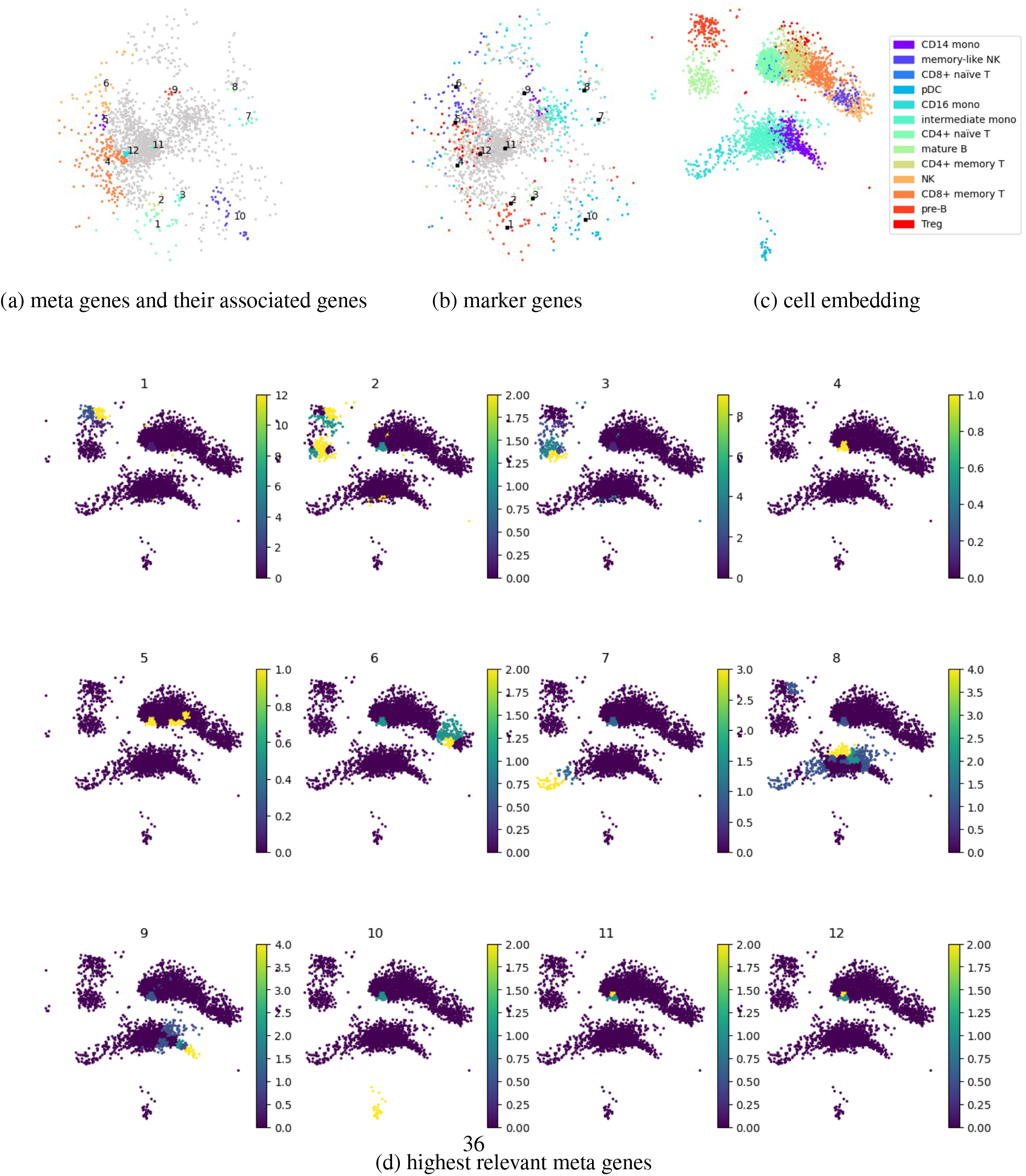
Exploration of the PBMC 5k (CITE-seq) dataset using a gene relevance map, which automatically detects correspondence between groups of cells and genes: (a) Gene embedding colored by genes associated with each meta-gene. (b) Gene embedding colored by marker genes specific to each cell type. (C) Cell embedding colored by cell types. (d) Gene relevance plot highlighting areas where the contribution of genes is highest. For example, meta-gene 10 exhibits high relevance in the lower region of the cell embedding.

**Figure S10:**
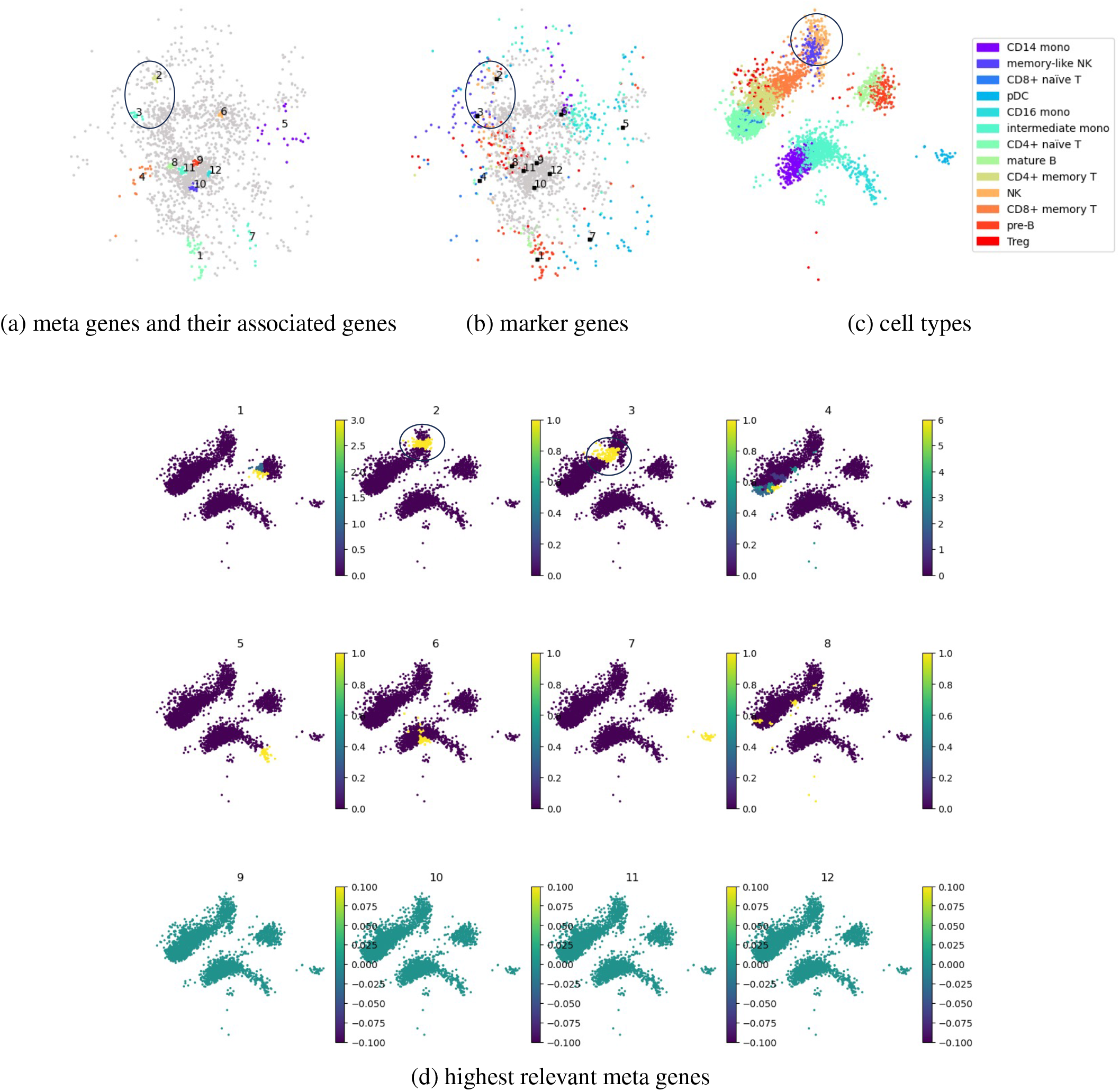
Exploration of the PBMC 5k (CITE-seq) dataset (multi-view version) using a gene relevance map, which automatically detects correspondence between groups of cells and genes: (a) Gene embedding colored by genes associated with each meta-gene. (b) Gene embedding colored by marker genes specific to each cell type. (c) Cell embedding colored by cell types. (d) Gene relevance plot highlighting areas where the contribution of genes is highest. For example, genes associated with meta-genes 2 and 3 are highly relevant to cells classified as NK and memory-like NK cell types, as indicated by the circles.

**Figure S11:**
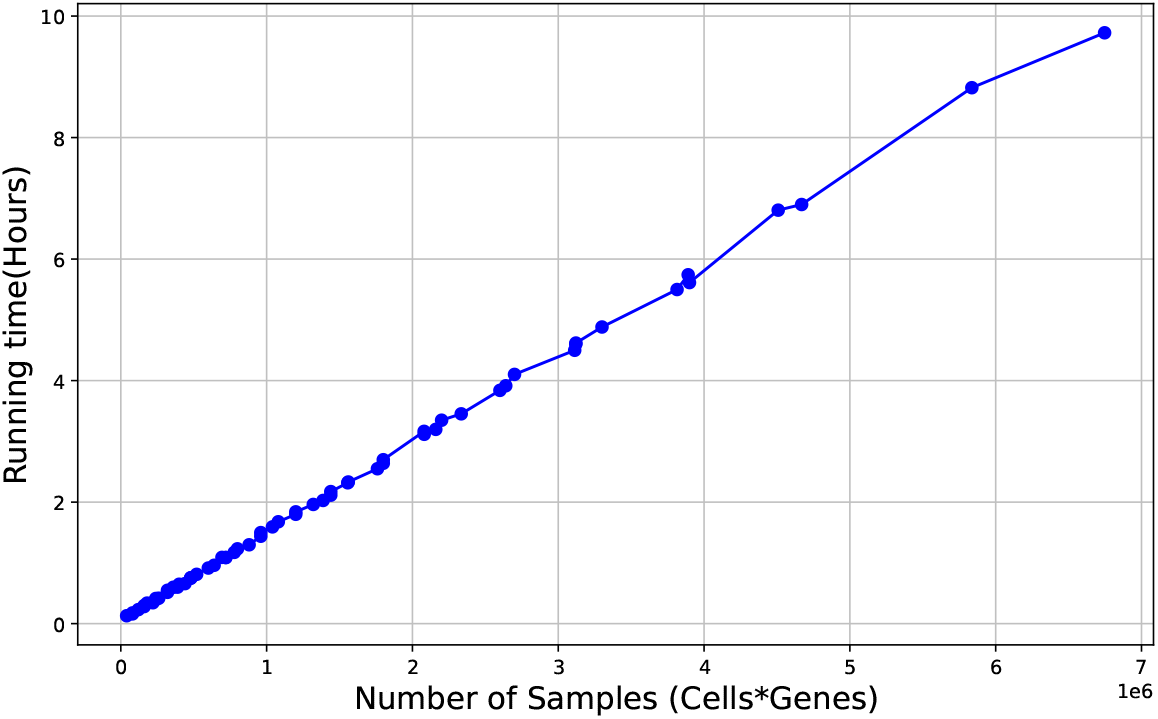
Running time of single-view MOMO-GP by increasing the size of observed data.

## 13 SUPPLEMENTARY TABLES

**Table S1:**
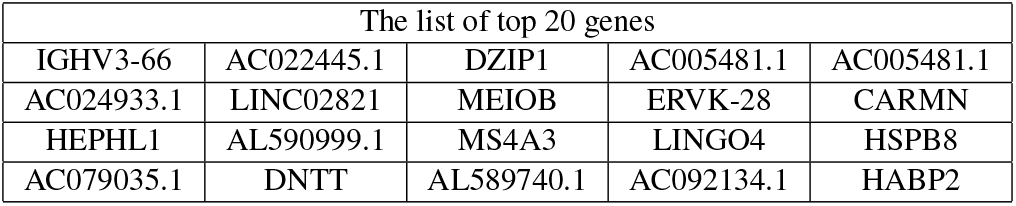
PBMC 10k dataset (scRNA-seq data): The list of top 20 genes located near the center of data embedded by MOMO-GP into 2D space.

**Table S2:** PBMC 5k (CITE-seq) dataset: A list of gene sets enriched for each meta-gene.

| Meta Gene | Term | Adjusted P-value | Combined Score | Cell-type Coverage |
| --- | --- | --- | --- | --- |
| 1 | Follicular B | 6.47-15 | 926.91 | 81.13 |
| 2 | Follicular B | 1.3e-08 | 1829.58 | 77.35 |
| 3 | Plasma | 6.76e-03 | 302.32 | NA |
| 3 | Follicular B | 1.01e-02 | 129.68 | 100 |
| 4 | Naïve T | 9.71e-82 | 3120.82 | 99.58 |
| 5 | NK | 8.4e-5 | 280.38 | 0.00 |
| 6 | NK | 4.77e-30 | 3560.08 | 100 |
| 7 | Monocyte | 4.72e-10 | 3685.82 | 100 |
| 9 | Neutrophil | 6.4e-5 | 382.07 | NA |
| 10 | Dendritic | 4.49e-16 | 1289.41 | 100 |

## Notes

### Competing Interest Statement

Florian Buettner reports funding from Merck KGaA and Bayer AG and renumeration from Albireo and Siemens AG

